# RdRpCATCH: A unified resource for RNA virus discovery using viral RNA-dependent RNA polymerase profile Hidden Markov models

**DOI:** 10.64898/2026.02.05.703936

**Authors:** Dimitris Karapliafis, Uri Neri, Ingrida Olendraite, Justine Charon, Shoichi Sakaguchi, Xin Hou, Dick de Ridder, Mark P. Zwart, Anne Kupczok

## Abstract

Recent advances in large-scale sequence mining have expanded our knowledge of RNA virus diversity. Most genome mining approaches for detecting RNA viruses that encode RNA-dependent RNA polymerase (RdRp) rely on identifying this conserved protein by employing profile Hidden Markov Models (pHMMs) to scan sequencing datasets. Recently, several new pHMM databases for RdRp detection have been released, each following distinct design principles. However, their relative performance is unclear and their accessibility to users without specialized computational expertise is limited. Here, we introduce the RdRp Collaborative Analysis Tool with Collections of pHMMs (RdRpCATCH: https://github.com/dimitris-karapliafis/RdRpCATCH), developed to consolidate publicly available RdRp pHMM resources into a single, accessible platform. RdRpCATCH enables the scanning of (meta)transcriptomic assemblies to discover RNA viruses and provides subsequent taxonomic annotation of detected contigs. A comparative analysis of RdRp pHMM databases reveals that most are highly effective at detecting known diversity of RNA viruses while minimizing false positives, supporting their joint use within RdRpCATCH. RdRpCATCH is distributed as both a conda package and a web server application (https://rdrpcatch.bioinformatics.nl), facilitating access for researchers with diverse expertise. By integrating multiple pHMM resources, this unified framework addresses fragmentation in the field and reduces technical barriers, enabling comprehensive viral discovery.

## Introduction

Viruses are the most numerous and diverse group of biological entities on Earth, with an estimated 10³¹ virus particles present on our planet (1, 2). Viral genomes consist either of DNA or RNA and have single or double-stranded conformations (3). Viruses with RNA genomes are characterized by high mutation rates, ranging from 10^−6^ to 10^−4^ mutations per nucleotide per round of replication, which is two orders of magnitude higher than DNA viruses (10^−8^ to 10^−6^ mutations per nucleotide per round of replication) and up to a million times higher than their cellular hosts (4, 5). These high rates, driven by rapid replication, large population sizes, error-prone replication mechanisms, and often the lack of proofreading domains that correct replication errors (6, 7), have led to an exceptionally diverse catalog of RNA virus species that has expanded greatly in recent years (8). This extensive diversity complicates the accurate detection of divergent viruses.

Traditional approaches for discovering and characterizing RNA viruses have relied on isolating and propagating infectious viruses in the laboratory, a process that is labor-intensive, time-consuming, and often limited by the availability of culturable host cells or organisms. Advances in high-throughput sequencing technologies over the past decade have enabled the study of RNA virus diversity at an unprecedented scale, revealing the ubiquity and diversity of RNA viruses across nearly all examined ecosystems (see Nagagawa et al. (9) for an overview). The International Committee for the Taxonomy of Viruses (ICTV) has formally recognized the impact of metagenomics by approving the use of metagenome sequence analysis for identifying new taxa (10). Despite these advances, current estimates suggest that less than 1% of existing RNA virus diversity has been identified (11, 12). Thus, the discovery of novel RNA viruses remains challenging due to their extensive genetic diversity, necessitating highly optimized computational detection methods tailored to their diversity.

*Riboviria* is a viral realm that encompasses all viruses encoding RNA-dependent polymerases -either RNA-dependent RNA polymerase (RdRp) or reverse transcriptase-for replication. Within *Riboviria*, the kingdom *Orthornavirae* comprises exclusively RNA viruses that encode an RdRp for genome replication, making RdRp the universal “hallmark” gene of this group. The core RdRp domain, approximately 230 amino acids in length, is structurally conserved and comprises three subdomains, called “palm”, “thumb”, and “fingers”, that collectively resemble a right-hand configuration (13). In contrast, sequence conservation is lower, especially across distant taxa, with some divergent RdRps sharing as little as 10% amino acid similarity (14). As a result, traditional sequence similarity-based methods often fail to detect highly divergent viral sequences, underscoring the need for more sensitive computational approaches. Profile Hidden Markov Models (pHMMs), probabilistic models generated by multiple sequence alignments of homologous sequences, have proven effective for remote homology detection, offering greater sensitivity than pairwise sequence comparison methods (15). Recently, deep learning-based approaches emerged for recognizing sequence or structural patterns in viral sequences (16–18). While these methods may enable detection of fainter homology signals than pHMMs, their validation and wider adoption remain limited, possibly due to the relatively high costs of protein structure prediction. In this regard, pHMM methods are still considered a balanced approach - being able to integrate the evolutionary fingerprint of proteins, allowing for deeper homology signal detection compared to primary sequence comparison-based methods such as BLAST (19) and performing on par with advanced deep learning methods (16).

Over the past five years, advances in sequencing technologies and genome mining have greatly expanded our understanding of RNA virus diversity. Multiple landmark studies have contributed significant discoveries. Wolf et al. (20) identified numerous RdRps in a metatranscriptome enriched for unicellular eukaryotes; the Serratus project (21, 22) and Neri et al. (23) both reported the detection of vast numbers of novel viruses at species-level operational taxonomic units across ecological samples and metatranscriptomes; the Tara Oceans project (24) revealed new phyla and classes of RNA viruses in marine samples; Olendraite et al. (25) highlighted RNA virus diversity in and beyond invertebrate hosts; and Hou & He et al. (16) leveraged deep learning to uncover remarkable viral diversity across diverse environments. Collectively, these efforts underscore the extensive scope of RNA virus diversity now being revealed. The widespread use of pHMMs for viral RdRp detection is reflected in the development of standalone tools and databases, such as the RdRp-scan pipeline -a curated and taxonomically labeled set of RdRp pHMMs and a sequence database designed to identify remote RdRp sequences in metatranscriptomic samples (26). Similarly, NeoRdRp 2.1 is designed to retrieve RdRps from metatranscriptomic datasets. Still, it differs in its construction strategy, relying on an extensive set of seed sequences and automated clustering to generate a comprehensive suite of pHMMs (27).

Most of the aforementioned studies have developed specialized pHMM databases targeting the RdRp sequence for virus detection, each employing distinct methodological approaches and seed sequences. The parallel development and their different design principles make it difficult to determine which database delivers superior performance for a given setting. Furthermore, technical challenges arise as databases are distributed across different repositories, and any effort to combine them into a single analysis pipeline requires users to manage multiple resources and is time-consuming. Using these databases separately and combining their results entails significant overhead in the “post-processing” of the results, as the different outputs may overlap substantially, increasing redundancy. This approach also directly affects how the statistical interpretation is conducted (e.g., multiple independent searches), as each database may pull in different types of false positives (FPs). To address this fragmentation and associated technical barriers, the RdRp Collaborative Analysis Tool with Collections of pHMMs (RdRpCATCH) was developed as a community initiative following the 1st RdRp Summit (28). RdRpCATCH consolidates all available RdRp pHMM databases into a unified resource, enabling users to scan metatranscriptomic assemblies, perform provisional taxonomic annotation of RdRp candidate sequences, and integrate custom RdRp pHMM databases. The tool is accessible as both a Python package (https://github.com/dimitris-karapliafis/RdRpCATCH) and a conda package, and through a web application (https://rdrpcatch.bioinformatics.nl).

## Materials & Methods

### Retrieval of pHMM databases

We retrieved the RdRp pHMM databases from their respective online resources (Table S1).

(i) LucaProt_HMM: This database was constructed through clustering and iterative search from 10,487 metatranscriptome assemblies (16). Starting with an initial set of 5,979 well-curated RdRp references from the RefSeq NCBI database, the assemblies were searched for RdRp matches. False positives were removed, leaving 713 RdRp clusters that were added to the query database. This refined database was used for subsequent searches, and the process was repeated 10 times, yielding a total of 754 pHMMs.
(ii) NeoRdRp: This database comprises 1,182 pHMMs constructed by sequences derived from the NCBI Virus database and a backbone set of RdRp-containing sequences (29).
(iii) NeoRdRp 2.1: NeoRdRp 2.1 expands the NeoRdRp collection to 19,394 profiles based on 565,093 sequences from various studies, which are clustered based on an automated pipeline (27).
(iv) Olendraite_fam: The Olendraite_fam database was constructed by aligning seed sequences from eukaryotic RNA viruses identified via GraviTy analysis (30) for family-level sequence-driven taxonomy, supplemented with RdRp sequences from NCBI nucleotide databases (25). This process involved 1,793 manually curated non-redundant core-RdRp sequences, yielding 77 family-level pHMMs.
(v) Olendraite_gen: The Olendraite genus-level database was generated from 10,912 RdRp sequences obtained from a previous study, Uniref-100, and by querying the TSA; genus-specific alignments then facilitated the construction of 341 pHMMs (31).
(vi) RdRp-Scan: This database was assembled from RdRp sequences retrieved from the NCBI non-redundant protein (nr) database, enriched with sequences from two viral metatranscriptomic studies and PalmDB (26). These sequences were aligned and manually curated to remove partial RdRp core sequences, resulting in 68 phylum- and order-specific pHMM profiles.
(vii) RVMT: This resource was constructed by analyzing 5,150 pre-assembled metatranscriptomes from IMG/M, which were then queried using tools such as psi-blast, hmmsearch, Diamond, and MMseqs2, utilizing seed sequences from prior research as initial profiles (23). Matches were filtered, trimmed to approximate the domain, clustered, and their alignments were used to identify additional sequences; this process was repeated twice before all identified RdRps were iteratively clustered and used in downstream analysis. Of these, 710 alignments for tentative families, orders, and classes were fetched and converted into pHMMs.
(vii) Zayed_HMM: An iterative process was employed to build this database, using a compilation of RdRp alignments from 66 virus lineages, approximately at the genus-family level, to generate the initial HMM database (24). These profiles were used to iteratively search 771 metatranscriptome assemblies derived from the Tara Oceans projects and other resources over 10 iterative cycles. Per cycle, the newly detected sequences were used to generate profiles, which were finally combined to yield 2,489 pHMMs.

Each database was formatted as an HMMER-pressed database using the hmmpress command of HMMER v3.3.2 (32). The pre-compiled databases are stored in Zenodo (https://doi.org/10.5281/zenodo.15463729).

The comparative analysis incorporated 12 pHMM profiles from Pfam-A v37.4 (33) that correspond to viral RdRps (RdRp_Pfam) as a baseline. Pfam RdRp profiles were selected as a reference point of accessible, curated, and community-accepted profiles, which have been used before for RNA virus discovery and detection (34, 35). These are not included in RdRpCATCH. The profiles were excluded because they exhibited lower sensitivity than the integrated databases in the subsequent comparative analysis. To construct this RdRp_Pfam dataset, we searched InterProScan and Pfam using the keyword ‘RdRp’ (36). This initial search produced 20 Pfam profiles, some of which are also found in cellular organisms. To focus on viral RdRps, we filtered out non-viral entries using the Pfam profile description. After this filtering step, we obtained a refined subset of 12 profiles: PF20489.3 (Birna_RdRp_C), PF04197.17 (Birna_RdRp_palm), PF04196.17 (Bunya_RdRp), PF22213.1 (CPV_RdRP_C), PF22209.1 (CPV_RdRP_N), PF22212.1 (CPV_RdRP_pol_dom), PF22152.1 (Permu_RdRp_palm), PF22260.1 (Permu_RdRp_thumb), PF00680.25 (RdRP_1), PF00978.26 (RdRP_2), PF00998.28 (RdRP_3), PF02123.21 (RdRP_4), and PF07925.16 (RdRP_5).These profiles were pressed to a pHMM database with the hmmpress command.

### RdRpCATCH implementation

RdRpCATCH is written in Python 3.12 and uses the pyHMMER3 v0.11.0 library for efficient pHMM searches (37). Interactive plots are generated using Altair v5.5.0 (38).

#### 1. Pre-processing of input sequences

For nucleotide sequences, length filtering is performed using Seqkit seq v2.10.0 (default: > 400 nt), followed by translation of contigs into six open reading frames with Seqkit translate v2.10.0, using the standard genetic code as a default (39). Stop codons are replaced by ‘X’ characters (an ambiguity character for HMMER) to preserve the genomic context of the query contig, even when a stop codon is identified under a specific genetic code.

#### 2. Hmmsearch-based RdRp detection

RdRp domains are identified using PyHMMER hmmsearch against multiple curated RdRp profile databases. Query sequences are searched against selected databases, and all hits meeting the specified E-value thresholds are retained for downstream processing.

#### 3. Definition of the RdRp hit coordinates

For each target sequence-profile pair, envelope coordinates (env_from, env_to) that define the probabilistic boundaries of domain matches are extracted from the hmmsearch output (32). When multiple domains of the same profile are detected within a single sequence, all domain envelope coordinates are aggregated to determine the boundaries of the RdRp region. The final RdRp start coordinate is the minimum env_from value across all detected domains, while the RdRp end coordinate is the maximum env_to value, creating a delimited region that encompasses all RdRp domain instances detected for that profile-sequence pair.

#### 4. Calculation of custom scores

RdRpCATCH calculates a series of custom scores that facilitate the evaluation of individual outputs and provide an overview of database performance across different datasets. To accurately quantify coverage in multi-domain hits, coverage metrics are determined for each target-profile pair using a custom interval-merging algorithm. This algorithm prevents double-counting of overlapping regions and ensures that gaps between domains are appropriately accounted for. For sequences containing a single detected domain, coverage is calculated as the interval length (env_from-env_to). In cases of multi-domain hits, all domain intervals are sorted by their start positions and iteratively merged: any overlapping or adjacent intervals (defined as intervals separated by ≥ 1 position) are combined, and the total coverage length is computed by summing the merged intervals.

In summary, the coverage scores calculated by RdRpCATCH are the following:

Profile Coverage: Fraction of the profile that aligns with the provided sequence, giving insight into how well the sequence matches the expected profile. Profile coverage is calculated as the merged length of all hmm_from and hmm_to intervals divided by the HMM profile length.

Sequence Coverage: Fraction of the sequence that aligns with the profile, helping assess the alignment’s completeness (assembled contig or protein sequence). Sequence coverage is calculated as the merged length of all env_from and env_to intervals divided by the target sequence length.

In addition, the following scores are calculated to enable fair comparison of sequences and profiles of varying lengths:

Normalized Bitscore (profile): The bitscore divided by the length of the profile. Normalized Bitscore (sequence): Bitscore divided by the length of the contig.

ID Score: Bitscore divided by the alignment length. Alignment length is calculated as the sum of the ali_from and ali_to intervals, corresponding to the actual aligned positions (25).

#### 5. Hit consolidation based on the best hit selection strategy

Following custom score calculation and coordinate determination for all target-profile pairs, a hierarchical best hit selection strategy is applied to identify the most confident RdRp hit for each sequence. First, within each profile database, the highest-scoring hit is selected for each target sequence, along with the total number of positive profiles detected in that database. Results from all searched databases are then consolidated. For sequences detected by multiple databases, the hit with the maximum bitscore across all databases is selected as the final best hit. Additionally, a summary annotation is generated for each sequence, indicating the total number of databases and profiles that detected the sequence, formatted as “database_name=number_of_profiles” and concatenated with semicolons for multiple databases.

#### 6. Taxonomic annotation of positive sequences

RdRpCATCH integrates provisional taxonomic assignment through MMseqs2 v17.b804f easy-taxonomy and easy-search modules against a custom RefSeq *Riboviria* database (40). To construct the database, protein sequences from RNA viruses were downloaded from the NCBI Virus portal (*Riboviria*, taxid:2559587; RefSeq only, downloaded on 11-02-2025), yielding 26,021 protein sequences. The dataset was formatted into a taxonomically informed MMseqs2 database using the NCBI taxdump and native MMseqs2 utilities, to retain the curated NCBI Taxonomy identifiers and lineage information already associated with each RefSeq entry, without performing any additional automatic taxonomic classification at this stage (41). NCBI RefSeq and NCBI Taxonomy were adopted as a single, internally consistent framework for sequence retrieval and taxonomic annotation in this study. MMseqs2 easy-taxonomy assigns taxonomic labels based on an accelerated approximation of the 2bLCA protocol (42).

### Reference dataset construction for comparative analysis

We retrieved previously detected RdRp amino acid sequence datasets and non-RdRp sequences from the literature (Tables S2, S3). As ground-truth RdRp-positive sequences, we collected:

(i) 161,979 RdRp representatives derived from 10,487 metatranscriptomes (16).
(ii) 77,510 full-length core RdRp domain sequences derived from 5,150 metatranscriptomes (23).
(iii) 12,110 ORFs containing RdRps from the Transcriptome Shotgun Assembly database (25).
(iv) 513,176 RdRp palmprint sequences clustered at 90% amino acid identity derived from the SRA database and various other resources (21).
(v) 7,080 core representative RdRp sequences from a previous study, derived from the NCBI Genbank, NCBI RefSeq, and a backbone RdRp set (43). These RdRPs were taxonomically assigned to 24 RNA virus “superclades” based on broad phylogenetic relationships, and subsequently to 5 phyla (out of seven total), 20 classes (23), 28 orders (33), and 112 families (139), based on both phylogenetic relationships and the International Committee on Taxonomy of Viruses (ICTV) taxonomy.
(vi) 4,593 RdRp sequences derived from a Yangshan Deep-Water Harbour metatranscriptome (20).
(vii) 6,238 core RdRp sequences derived from 771 Tara Oceans metatranscriptomes (24).

We collected two non-RdRp datasets as true-negative datasets:

(i) The dataset denoted as Lucaprot_TN is a negative dataset used in model training from the LucaProt study, which comprised a total of 229,434 sequences. Specifically, it contains eukaryotic RNA-dependent RNA polymerases (Eu RdRP, N = 2,233), eukaryotic DNA-dependent RNA polymerases (Eu DdRP, N = 1,184), reverse transcriptases (RT, N = 48,490), proteins obtained from DNA viruses (N = 1,533), non-RdRp proteins obtained from RNA viruses (N = 1,574), and a wide array of cellular proteins from different functional categories (N = 174,420).
(ii) The custom set of true negative sequences, denoted as custom_TN, comprises 576,624 sequences and were retrieved from Swissprot (44) using the following queries: 20,211 polymerases that are not viral, with the query (id:Polymerase) OR

(protein_name:Polymerase) OR (cc_function:Polymerase) OR (ec:2.7.7.-) NOT (taxonomy_id:2732396) NOT (taxonomy_id:2788808); 544,276 cellular proteins retrieved using the query: NOT (taxonomy_id:2559587) NOT (taxonomy_id:10239) NOT (cc_function:Polymerase); and 12,137 non-*Orthornavirae* viral proteins retrieved with the query (taxonomy_id:10239) NOT (taxonomy_id:2732396).

### Comparison of RdRp pHMM database performance

To evaluate the performance of the databases, we designed two distinct comparative analysis scenarios, each using different combinations of the aforementioned datasets.

(i) Comparative analysis 1: In the first scenario, we explored the utility of the databases in detecting a well-defined set by using the 7,080 VirID_core_RdRps (mean length: ∼1712 aa, Fig. S9). As a ground-truth negative dataset, we used Lucaprot_TN, which comprises a diverse collection of 229,434 cellular and viral non-RdRp sequences (mean length: ∼450 aa, Fig. S9).
(ii) Comparative analysis 2: In the second scenario, we aimed to test the utility of the databases in retrieving literature-retrieved diverse RdRp sequences. For this purpose, we aggregated all available RdRp amino acid sequence datasets into a single file, resulting in a combined set of 782,686 RdRp sequences. These sequences were then clustered at 90% sequence identity and 70% coverage using MMseqs2 v17.b804f, with flags --cov-mode 1 --cluster-mode 2 (greedy incremental clustering). Under these settings, the longest sequence in the cluster becomes the representative sequence. This analysis yielded a consolidated ground truth dataset of 525,244 RdRp representative sequences (mean length: ∼212 aa, Fig. S9). The length of the sequences in this dataset is considerably smaller than for VirID_core_RdRps only, due to the integration of trimmed-to-palmprint RdRp sequences from PalmDB. For the negative set, the Lucaprot_TN dataset was merged with the custom_TN set of 576,624 non-RdRp sequences retrieved from SwissProt. Both negative datasets were clustered with MMseqs2 v17.b804f (using the same settings as for the true positives), yielding a dataset of 560,209 dereplicated true-negative sequences (mean length: ∼409 aa; Fig. S9).

A Python script using PyHMMER hmmsearch was used to scan the true-positive and true-negative datasets independently with each database. Hmmsearch was configured with the following parameters: (i) the E-value threshold was set to 1, which was the most permissive threshold included in the analysis; and (ii) the effective number of sequences (Z) was fixed at 1,000,000 to provide a fair comparison across different datasets. Subsequently, a series of E-value thresholds (10^0^, 10^−1^, 10^−2^, …, 10^−15^) was applied iteratively to the raw hmmsearch output. For each tested threshold, the corresponding true positives (TP), true negatives (TN), false positives (FP), and false negatives (FN) were calculated per database. We also evaluated performance by combining the results from all databases, consolidating positive and negative hits, and assigning the ‘All_databases’ flag to these combined results. This approach provides an overall assessment of detection sensitivity and specificity when leveraging all databases together. Sequences from the positive dataset detected by any database were counted as TPs, while those not detected by any database were FNs.

Sequences from the negative dataset detected by any database were FPs, and those not detected by any database were TNs. Using these, the following performance metrics were derived:

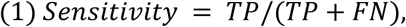

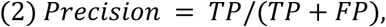

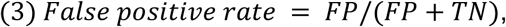

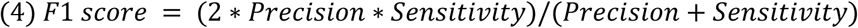

To investigate the taxonomic distribution of false-positive sequences identified by the databases, we extracted UniProt Identifiers from the headers of false-positive sequences detected at distinct E-value thresholds. Subsequently, we used these identifiers as input to the ID-mapping module of the UniProt web interface (45). The ID-mapping file, which contained the taxonomic annotations for the input sequences, was downloaded and used to map Uniprot IDs to HMM hits. For taxonomic assignment, it was first checked if the entry was viral by searching for the term “Viruses” within the lineage string. If identified as viral, it was further sub-classified by its viral kingdom (e.g., *Orthornavirae*). If not viral, the script assigned the sequence to a cellular domain (Bacteria, Eukaryota, or Archaea) based on the highest-level domain present in its lineage. Entries noted as “deleted” in UniProt or that could not be mapped were excluded from the analysis (573 and 574 sequences for the first and second comparison scenarios, respectively). To compare the taxonomic distribution of FP hits across the different HMM databases, the classified data were aggregated for each E-value threshold.

To assess whether bitscore thresholds can be employed in addition to E-values and help control false positives, we identified a bitscore cut-off that maximizes the F1 score (the harmonic mean of precision and sensitivity) for each examined E-value. The F1-score was selected as the primary optimization metric due to its robustness to imbalanced datasets (where one class has significantly fewer examples than others). For each E-value threshold, we iteratively scanned integer bitscore values ranging from 0 to the maximum bitscore detected per database, and calculated the F1 score for each of these bitscores. The bitscore that yielded the maximum F1 score was selected as the optimal cutoff for the given e-value. Subsequently, the maximum F1 score across E-values was chosen for each database; in cases where multiple E-value and bitscore threshold combinations resulted in an identical maximal F1 (evaluated to a precision of four decimal places), the combination associated with the least stringent (i.e., highest) E-value was selected as the single best-performing result for annotation purposes.

### Performance assessment on publicly available datasets

To evaluate the performance of RdRpCATCH on publicly available data, we reanalyzed data from a recent study investigating viral diversity in moss (46). This study included 16 *Sphagnum*-derived transcriptomes that were host-filtered and *de novo* co-assembled with MEGAHIT v1.2.9 (47). We downloaded the resulting co-assembly from Zenodo (https://zenodo.org/records/15281951) and filtered contigs for a minimum length of 2,000 nt. We applied geNomad v1.12 (48) in two configurations (default and conservative) to this dataset, enabling the score calibration module. RdRpCATCH was applied to the same dataset, in three configurations: RdRpCATCH-efficient (using RdRp-Scan and Olendraite_fam databases). RdRpCATCH-balanced (using Lucaprot_HMM, RVMT and Zayed_HMM) and RdRpCATCH-full (using all supported databases). For RdRpCATCH the E-value was set to 10^−5^ (default) and the flag --length-thr to 2,000. All contigs were submitted to InterProScan v5.77-108.0 (49) and the output was parsed for RdRp evidence: hits of the Pfam profiles described above as RdRp_Pfam, hits against non-Pfam RdRp InterProScan profiles (PS50507, PS50525, PS50526, PR00914) or the presence of RdRp-related substrings (i.e., rdrp, rna-dependent rna polymerase, rna-directed rna polymerase, rna directed rna polymerase, rna-directed rna-polymerase, rna dependent rna polymerase) were considered supporting of RdRp presence. Subsequently, all viral contigs were submitted to Palmscan v1 (21) (-hiconf) to validate the presence of the RdRp catalytic motifs A, B and C. Finally, all contigs were submitted to MMseqs2 v17.b804f easy-taxonomy search against Uniref100 (downloaded on 17-4-2026) and were stratified to the following categories: (i) Cellular, (ii) Unclassified sequences, (iii) Viral (*Orthornavirae*), (iv) Viral (*Riboviria*-unclassified incl. *Pararnavirae*) and (v) Viral-unclassified according to their taxonomic annotations to facilitate downstream comparison of the results.

All analyses were performed on a 128-core AMD EPYC 7532 system with 3 TB of RAM, using 20 threads.

## Results

The RdRp Collaborative Analysis Tool with Collections of pHMMs (RdRpCATCH) enables scanning of metatranscriptomic assemblies or protein sequences for viral RdRp signals using a pre-compiled collection of RdRp pHMMs. RdRpCATCH employs a modular architecture that facilitates the integration of multiple RdRp pHMM databases (Fig. 1). Users may also scan sequences with their own custom RdRp domain databases and combine these results with any selection of pre-compiled databases. Additionally, the modular architecture of RdRpCATCH allows for the integration of newly developed databases in the future. The software package provides automated utilities for downloading and setting up databases, facilitating access to and maintenance of current versions of all integrated pHMM databases. Command-line interface options support batch processing and integration into existing bioinformatics pipelines. Comprehensive logging and output formatting support reproducibility and facilitate downstream analysis.

**Figure 1.**
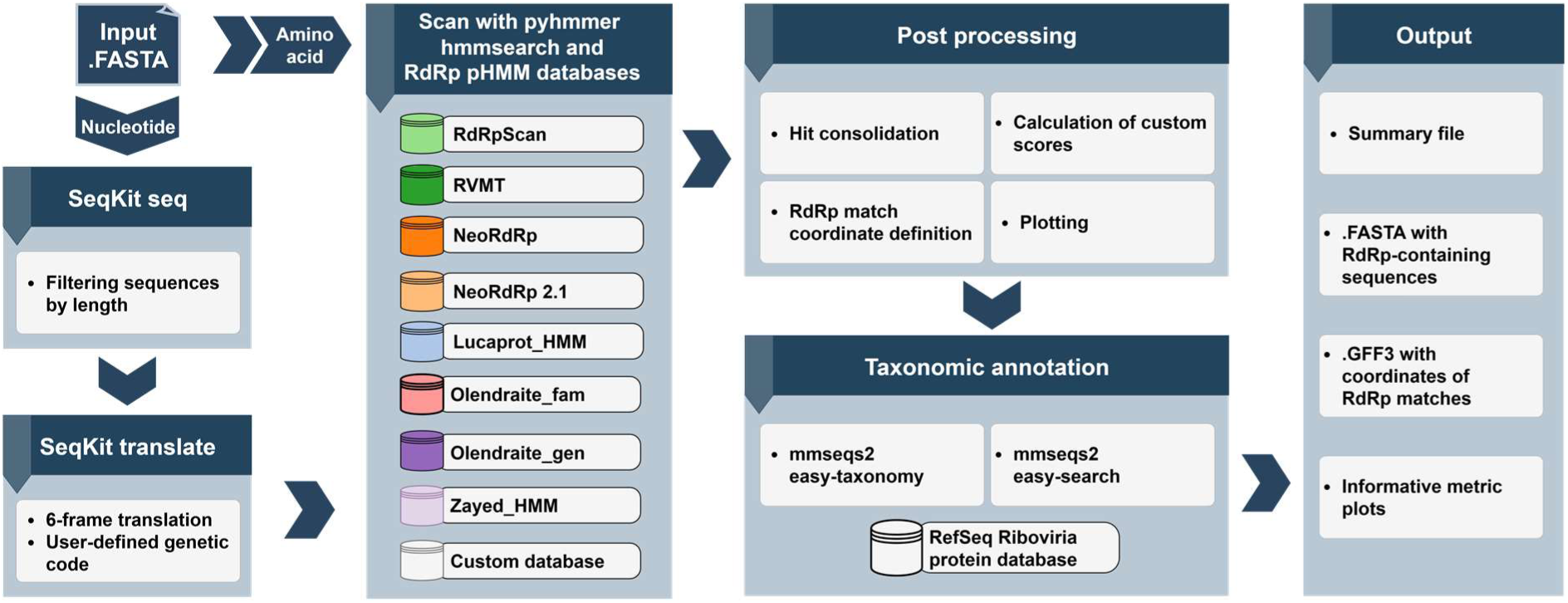
The RdRpCATCH workflow.

### RdRpCATCH workflow

RdRpCATCH integrates eight pHMM databases specifically developed for RdRp detection, collectively comprising 25,055 pHMM profiles (Fig. 2, Fig. S1). Each database was pre-compiled with its original architecture intact (no models were added or removed), preserving the unique optimization and performance characteristics intended by its creators. This approach leverages the full capabilities of each contributing database while offering a unified interface for comprehensive viral detection.

**Figure 2.**
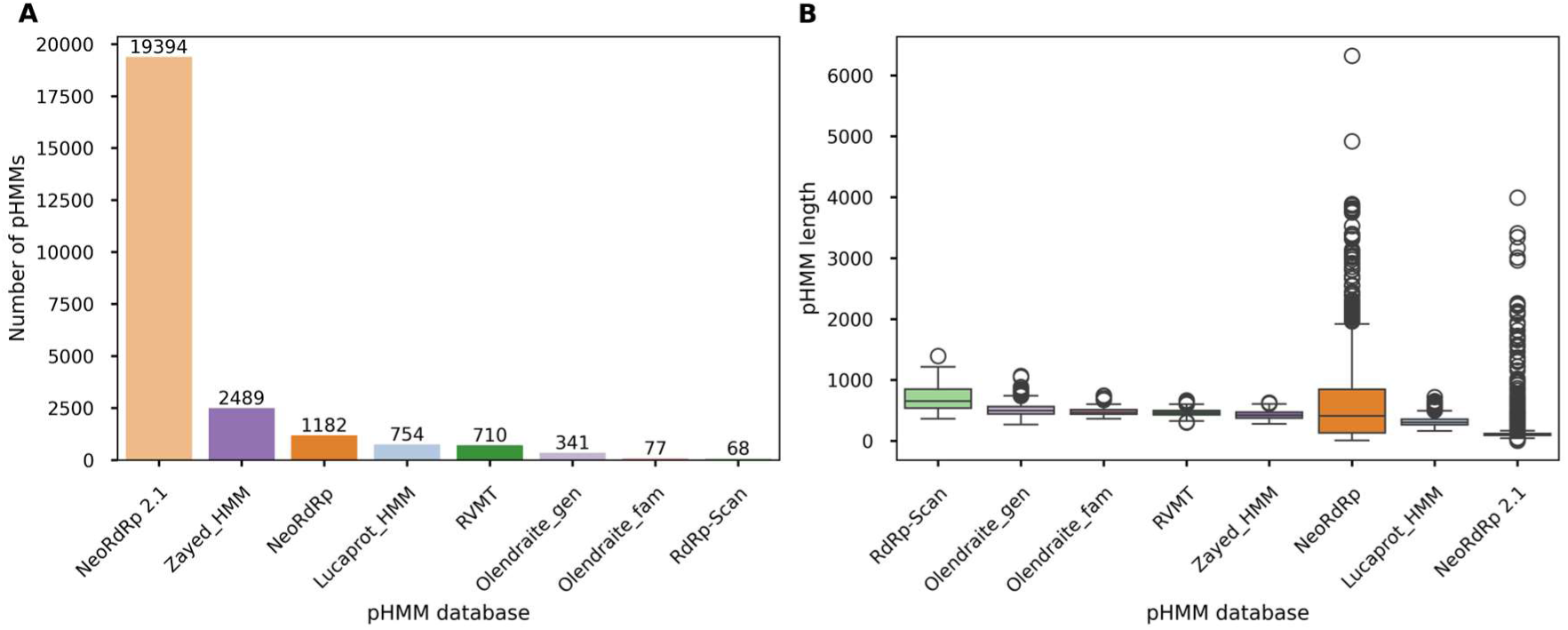
(A) Total number of profile HMMs per database. (B) Profile-HMM length distributions for different databases.

RdRpCATCH accepts input as FASTA files containing either amino acid or nucleotide sequences. Several RNA virus taxa utilize alternative genetic codes; for instance, in the family *Mitoviridae* (50). Consequently, the choice of translation method can influence the detection of such viruses in transcriptomic datasets. RdRpCATCH addresses this by performing a six-frame translation of complete contigs and supporting multiple genetic codes (as query options). In the next step, the sequences are scanned against the pre-compiled set of RdRp pHMM databases using PyHMMER hmmsearch. To minimize redundancy and provide streamlined results for downstream analysis when multiple RdRp profiles align, RdRpCATCH implements a best-hit selection algorithm that identifies the highest-scoring match across all integrated databases for each input sequence. The coordinates of the RdRp core domain sequence are determined based on the envelope coordinates of each hit. Based on the RdRp core domain coordinates, RdRpCATCH generates trimmed amino acid FASTA files, complete amino acid sequence FASTA files, and complete nucleotide sequence FASTA files when applicable. Additionally, RdRpCATCH outputs a GFF3 file containing RdRp domain coordinates on the amino acid sequence. Finally, the positive sequences are annotated taxonomically using MMseqs2 and a RefSeq *Riboviria* protein database, providing an efficient provisional taxonomic context for detected sequences.

### Software availability and web server

RdRpCATCH is distributed as a Bioconda package (rdrpcatch) and as a Python package on PyPI, enabling command-line use for scanning nucleotide or protein FASTA files. In addition, RdRpCATCH is available as a web server (https://rdrpcatch.bioinformatics.nl) that allows users to upload assemblies via a drag-and-drop interface, run RdRp pHMM searches against the supported databases and provide downloadable result tables and visualizations for further exploration of putative RNA virus contigs.

### Comparative analysis 1: Performance of integrated databases in RdRpCATCH using a well-defined set of RdRp sequences

To evaluate the performance of the supported databases in detecting viral RdRps, we designed two comparison scenarios. In the first experiment, we assessed how effectively the databases could identify a defined set of 7,080 RdRp sequences obtained from the VirID study that were annotated with the ICTV taxonomy. To determine their ability to differentiate between actual RdRp sequences and non-RdRp sequences, we also deployed the databases against a dataset of 229,434 non-RdRp sequences that had previously been used to train the LucaProt deep learning model (16).

F1 scores indicated that most supported databases outperformed RdRp_Pfam across all tested E-value thresholds (Fig. 3A). RVMT, Zayed_HMM (0.9997), and Lucaprot_HMM (0.9995) achieved the highest F1 scores at stringent E-value thresholds (Fig. 3F). RdRp-scan also performed strongly (0.9983) across a range of thresholds, while Olendraite_fam (0.9920) and Olendraite_gen (0.9885) improved as thresholds became less stringent. NeoRdRp 2.1 and NeoRdRp showed lower F1 scores, especially at relaxed thresholds (0.9475 and 0.8667, respectively). Overall, the F1 score declined rapidly (Zayed_HMM, RVMT, Lucaprot_HMM, NeoRdRp, NeoRdRp 2.1) or plateaued (RdRp-scan, Olendraite_gen, Olendraite_fam) after an E-value of 10^−5^, indicating that this threshold achieves a good trade-off between sensitivity and precision. Since the F1 score combines both precision and sensitivity, the following sections analyze each metric separately to clarify its individual contributions to overall performance.

**Figure 3.**
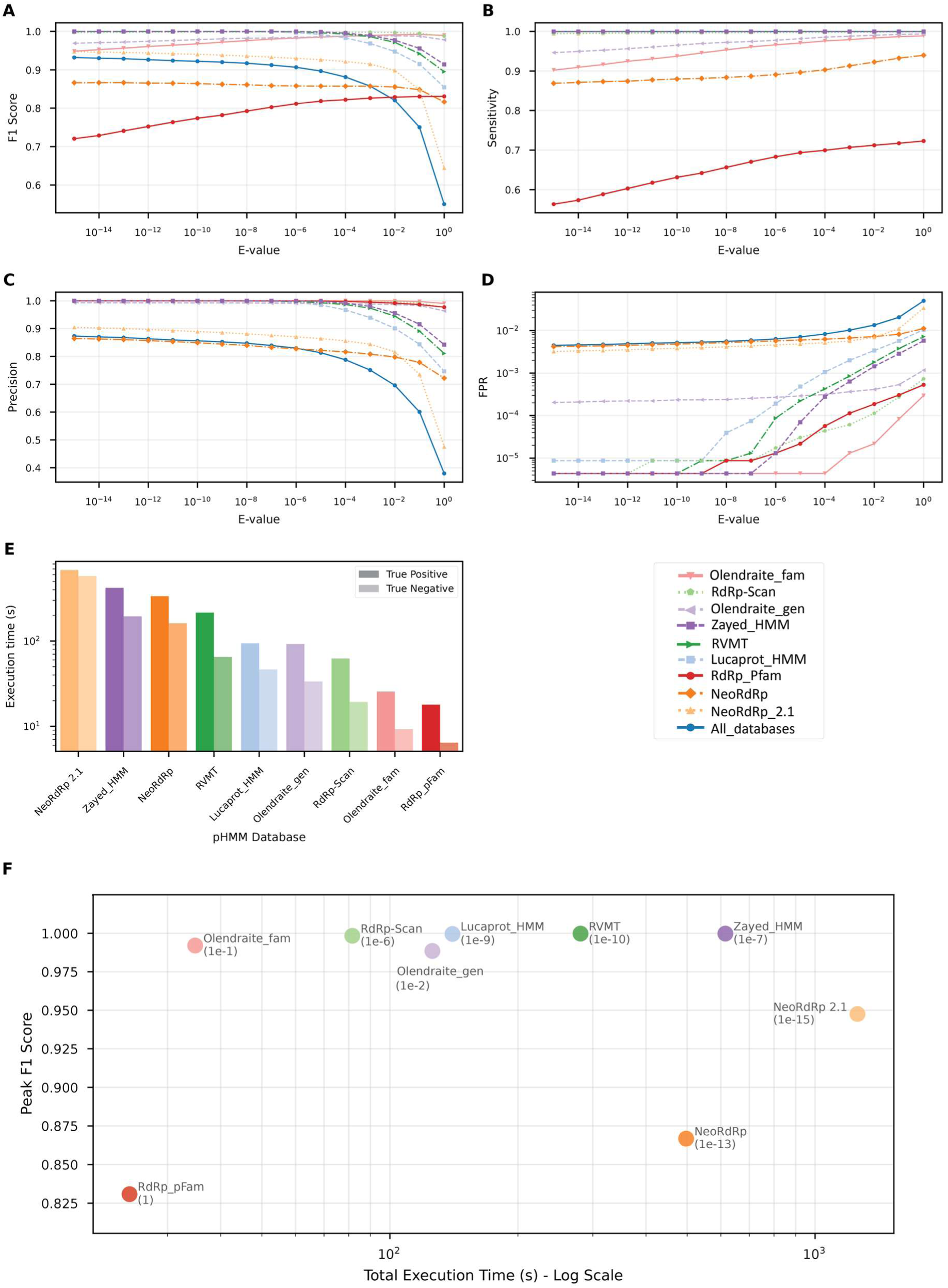
Performance of the RdRp HMM databases for comparative analysis 1 (VirID_TP/Lucaprot_TN datasets) across a range of E-value thresholds, from 10^−15^ to 1. (A) F1-Score: The harmonic mean of precision and sensitivity, providing a balanced measure of a database’s overall accuracy. (B) Sensitivity (Recall): The proportion of true positive sequences correctly identified (True Positive Rate). This plot shows each database’s ability to detect known RdRp sequences. (C) Precision: The proportion of positive identifications that were correct. This metric highlights a database’s ability to avoid false positives.(D) False Positive Rate (FPR): The proportion of negative sequences incorrectly identified as positive. The y-axis is log-scaled to better visualize differences at low error rates. (E) The execution time (on a logarithmic scale, in seconds) required to search against various RdRp HMM databases per dataset. (F) Peak F1 score vs. total runtime per database; labels show database name and optimal E-value.

Sensitivity reflects how effectively each database detects true positives, in this case the number of viral RdRp sequences in the ground truth dataset. For most databases, sensitivity increased with E-value, allowing detection of more distantly related sequences (Fig. 3B). RVMT, Zayed_HMM and Lucaprot_HMM achieved their highest sensitivity already at stringent thresholds (E<=10^−5^), each detecting 7,078 out of 7,080 sequences (0.9997). NeoRdRp 2.1 also reached the sensitivity of 0.9987 but for E = 1, while RdRp-scan peaked at 0.9987 (7071 sequences) at a similarly relaxed threshold (E = 0.1). Olendraite_gen (0.9935, 7034 sequences), Olendraite_fam (0.9888, 7001 sequences), and NeoRdRp (0.9396, 6653 sequences) reached their highest sensitivity at the most relaxed threshold tested (E = 1). Combining all databases boosts sensitivity further to 0.9998 (7079 sequences) at E = 0.1. Importantly, even the least sensitive databases outperformed the RdRp_Pfam reference (0.7228, 5118 sequences), and most achieved nearly identical peak sensitivities, indicating strong detection capabilities.

Precision, or the proportion of correctly identified positives, generally decreased as E-values were increased (Fig. 3C). RVMT, Zayed_HMM, Olendraite_fam, RdRp-scan, Lucaprot_HMM, and Olendraite_gen maintain high precision (>0.99) at stringent thresholds (E < 10⁻⁵), but precision for RVMT, Zayed_HMM, and Lucaprot_HMM declined at higher E-values. NeoRdRp and NeoRdRp 2.1 showed lower maximum precision and lost precision more rapidly as thresholds relaxed, indicating a comparatively higher FP rate. When combining all databases, overall precision dropped further due to cumulative FPs. This highlights the key trade-off: relaxing E-value thresholds increased sensitivity but reduced precision, and hence, the decline in F1 score is attributed to declining precision.

A low false positive rate (FPR) is essential for RNA virus discovery from environmental metatranscriptomic samples because most assembled contigs originate from cellular hosts rather than viruses. We assessed FPR across E-value thresholds and found that under strict settings (E ≤ 10^−6^), Olendraite_fam and RdRp-scan misclassified fewer than one in 200,000 sequences, with FPR remaining low until E-values increased (Fig. 3C). As thresholds relaxed, FPR rose for all databases: for example, Zayed_HMM, RVMT, and Lucaprot_HMM started at an FPR of around 1 in 200,000 for E = 10^−15^ but increased one-fold or two-fold at moderate thresholds. NeoRdRp variants had higher baseline FPRs and exceeded 1 per 100 sequences at the most relaxed settings. The FPR of the combined approach rose to nearly 4 in 100 at E = 1. To provide a qualitative overview of the false positive sequences, we analysed their taxonomic annotation and their distribution across E-values (Fig. 4). For strict E-values, most FPs found by NeoRdRp, NeoRdRp 2.1, and Olendraite_gen were of viral origin, often from the *Orthornavirae* or unclassified groups, suggesting that some profiles include non-RdRp domains. On the contrary, for higher E-values (mainly E > 0.01), FPs were dominated by cellular sequences across all databases, highlighting the need for filtering steps to maintain the validity of the analysis. Analysis of profile lengths, particularly for NeoRdRp and NeoRdRp 2.1, revealed outliers exceeding 1,500 amino acids, supporting the observation that the profiles may contain non-RdRp domains (Fig. 2).

**Figure 4.**
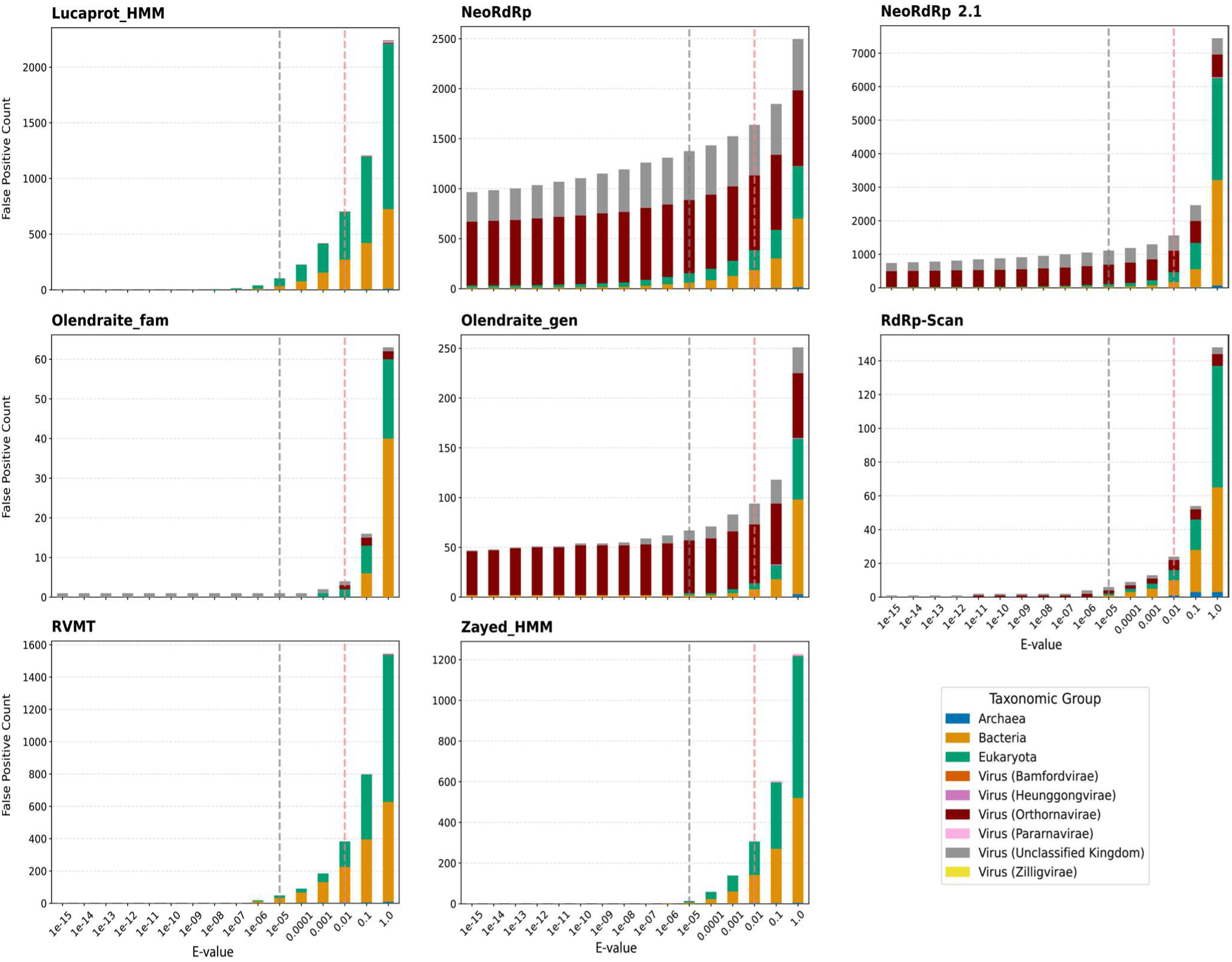
Taxonomic breakdown of false positive hits across E-value thresholds for the Lucaprot_TN dataset. Dashed lines indicate an E-value of 10^−5^ (grey) and 10^−2^(light red) to indicate recommended and default thresholds for hmmsearch.

Bitscores may provide a practical, database-independent method for controlling FPs. The distribution of FP bitscores shows that, at relaxed E-values, low-bit-score FPs are retrieved (Fig. S2). This pattern is especially evident for RdRp-scan, RVMT, Olendraite_fam, Lucaprot_HMM, and Zayed_HMM. To evaluate whether a bitscore threshold can filter FPs, we identified a bitscore cutoff that maximizes the F1 score for each examined E-value. For all databases that showed a decrease in F1 and an increase in FPR at high E-values, the effect was mitigated by employing a bitscore threshold (RdRp_pFam: 14, Olendraite_fam: 18, Olendraite_gen: 19, Zayed_HMM: 35-74, RdRp-scan: 36, RVMT: 44-54, Lucaprot_HMM: 44-58, NeoRdRp: 58; NeoRdRp 2.1: 98) (Figs. S3, S4). These results indicate that, regardless of the E-value, the databases can produce reliable results when supplemented with a bitscore filter as an additional safeguard. However, it is unclear whether these thresholds can be generalized to more complex datasets, which may include fragmented or novel RdRp sequences.

Finally, we assessed how database size and sequence volume affect runtime. The larger TN set was processed more quickly than the TP set because HMMER’s heuristic filters efficiently eliminate non-matching sequences (Fig. 3E). For true positives, runtime closely reflected the number of pHMMs and the length of the profiles in each database: the smaller databases, Olendraite_fam and RdRp-scan, finished in 18 and 62 seconds, respectively, whereas larger databases, such as NeoRdRp 2.1 (19,394 profiles), took up to 679 seconds. Intermediate-sized databases such as Olendraite_gen, Lucaprot_HMM, and RVMT showed intermediate runtimes (ranging from ∼92 to 335 seconds). Thus, runtime is primarily determined by database size, with larger databases requiring substantially more analysis time.

### Comparative analysis 2: Performance when investigating an extensive dataset of literature-retrieved diverse RdRps

In the second comparative analysis scenario, we used a much larger and more diverse dataset: 525,244 amino acid RdRp sequences from multiple studies (clustered at 90% sequence identity and 70% coverage) as positives, and 560,209 non-RdRp sequences compiled from merging the Lucaprot_TN dataset with a custom set of non-RdRp proteins retrieved from UniProt, using the same clustering thresholds (see Materials & Methods). This dataset presents a more challenging setting for the databases due to its composition of diverse RdRp sequences, with a significant portion trimmed to the small, conserved palmprint subdomain (∼250 aa).

Most databases achieved F1 scores above 0.95 (Fig. 5A), performing similarly to the first scenario and far surpassing RdRp_Pfam. Notably, for most databases, F1 scores remained stable or even improved at higher E-values, rather than declining. The combination of databases produced the highest overall F1 score (0.9962), while Lucaprot_HMM was the top-performing individual database (0.9954). RVMT, Zayed_HMM, NeoRdRp 2.1, and RdRp-scan also demonstrated strong results, particularly at more relaxed thresholds (Fig. 5F). Olendraite_gen, NeoRdRp, and Olendraite_fam consistently improved as the E-value threshold increased. Sensitivity trends closely mirrored those of the F1 score, highlighting the impact of this metric for the performance in this dataset (Fig. 5B). Precision, on the other hand, showed minimal variation: all databases maintained exceptionally high precision, with only slight declines at higher E-values (Fig. 5C). Unlike the first scenario -which featured a positive-to-negative sequence ratio of about 1:32-this dataset is nearly balanced between positive and negative sequences (∼0.94:1). Consequently, FPs and FNs carry comparable weight in this scenario, whereas in the first scenario, the impact of FPs in precision is inflated. However, environmental metatranscriptomic assemblies are more likely to resemble the first scenario, with cellular sequences expected to vastly outnumber viral RdRp sequences. Analysis of FPR trends revealed patterns very similar to those observed in the first scenario, with only slight improvement (Fig. 5D). Examining the taxonomic distribution of false positive sequences showed the same pattern as the initial setting, with most FPs assigned to viral sequences (with the majority belonging to *Orthornavirae* or unclassified kingdoms), for stringent E-values, from the NeoRdRp, NeoRdRp 2.1, and Olendraite_gen databases (Fig. S5). This further supports the hypothesis that some profiles may include additional regions of viral genomes, not just the RdRp domain.

**Figure 5.**
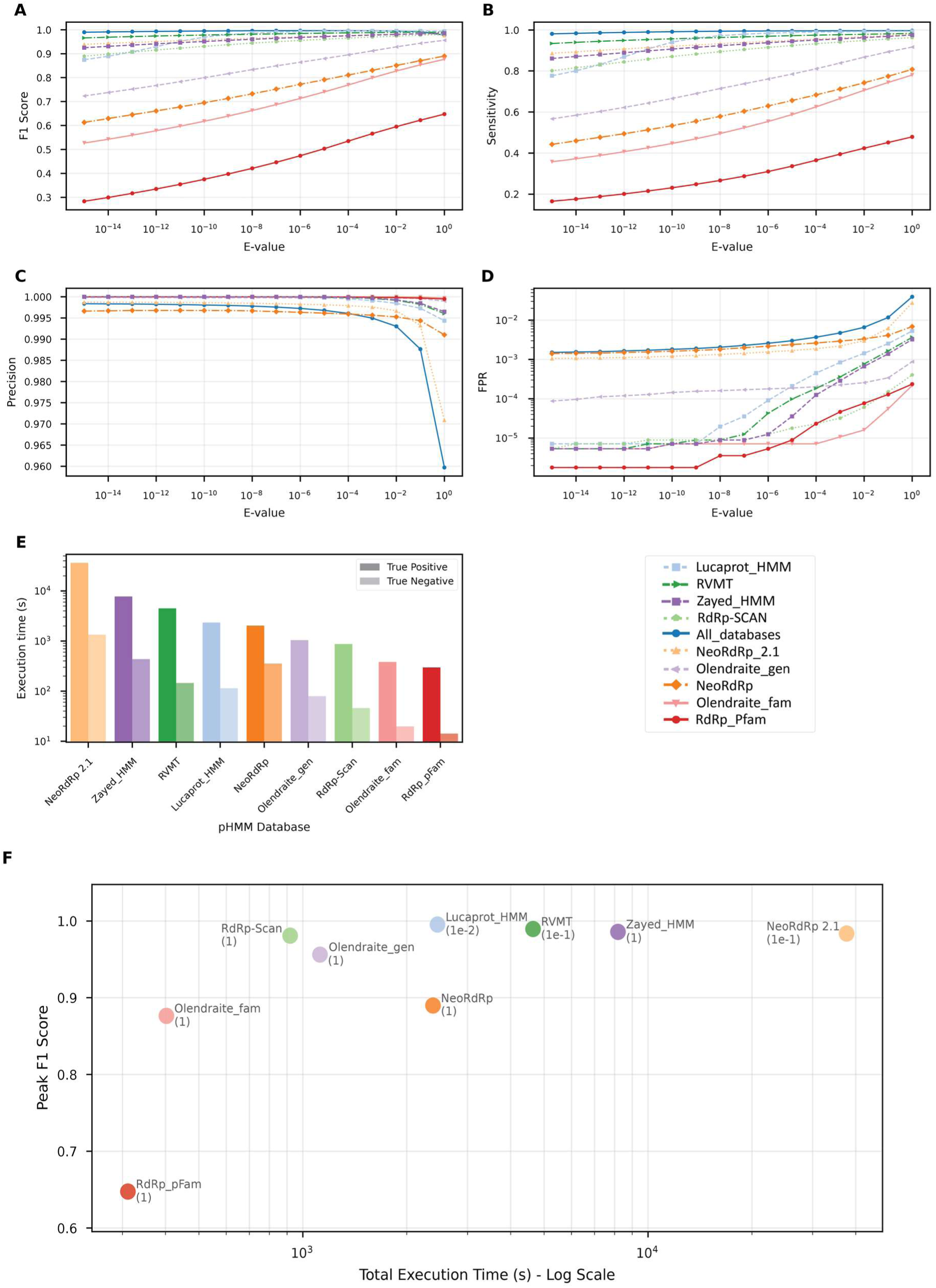
Performance of the RdRp HMM databases for comparative analysis 2 (Total_TP/Total_TN dataset) across a range of E-value thresholds, from 10^−15^ to 1. (A) F1-Score: The harmonic mean of precision and sensitivity, providing a balanced measure of a database’s overall accuracy. (B) Sensitivity (Recall): The proportion of true positive sequences correctly identified (True Positive Rate). This plot shows each database’s ability to detect known RdRp sequences. (C) Precision: The proportion of positive identifications that were correct. This metric highlights a database’s ability to avoid false positives. (D) False Positive Rate (FPR): The proportion of negative sequences incorrectly identified as positive. The y-axis is log-scaled to better visualize differences at low error rates. (E) The execution time (on a logarithmic scale, in seconds) required to search against various RdRp HMM databases per dataset. (F) Peak F1 score vs. total runtime per database; labels show database name and optimal E-value.

Analyzing the bitscore trends revealed a key difference between the two benchmarking scenarios: In the second scenario, the bitscore distributions for TPs and FPs overlap much more than in the first scenario (Fig. S6). This is primarily because the true-positive set here is more diverse, including many smaller RdRp palmprint sequences from PalmDB. It is observed that shorter sequences tend to exhibit lower bit scores for true positives (Fig. S10, S11). Therefore, there is a considerable number of short and/or divergent RdRp palmprint sequences that are detectable by pHMMs; however, even if they contain the most conserved part of the RdRp sequence, their bitscores are similar to those of high-scoring false positives. As a result, applying the optimal bit score thresholds did not improve the F1 score and FPR, as in the previous scenario, since filtering out FPs also removed a similar number of true positives (Fig. S7, S8). This suggests that bitscore thresholds optimized for one dataset may not generalize to more diverse or fragmented metatranscriptomic samples, where small or incomplete contigs are common but can still be detected by sensitive HMM profiles. In such cases, we suggest implementing robust downstream methodologies to control FPs in comprehensive discovery scenarios.

Finally, we compared runtimes for the two datasets (Fig. 5E). As before, the negative set was processed much faster than the positive set, due to HMMER’s ability to discard non-matching sequences quickly. For example, Olendraite_fam was processed in 20s versus 383s for the negative versus positive sets, respectively. Larger databases, such as NeoRdRp 2.1, took 327s for negatives and up to 36,297s for positives, demonstrating that positive-set runtimes scale sharply with sequence volume and database size and underscoring the importance of careful resource planning in large-scale analyses.

### Practical recommendations and database selection guidelines

Our comparative analyses reveal that database performance is significantly influenced by dataset diversity, the selection of search thresholds, and the availability of computational resources. Across both comparison scenarios, most databases achieved high F1 scores and maintained low false positive rates when stringent E-value thresholds were applied. In particular, Lucaprot_HMM, RVMT, and Zayed_HMM consistently provided the best overall balance between sensitivity, precision, and low FPR and are thus suggested for RdRpCATCH-balanced mode. For large-scale analyses, where minimizing FPs and achieving rapid runtimes are priorities, RdRp-scan and Olendraite_fam stood out and are thus suggested for RdRpCATCH-efficient mode. These databases offered high precision and fast processing speeds, making them well-suited for screening extensive sequence datasets (Fig. 3F, 5F). However, this efficiency came with a trade-off in sensitivity, particularly for Olendraite_fam, when compared to the top performers. NeoRdRp 2.1, on the other hand, demonstrated good sensitivity; however, this came at the cost of higher false positive rates and considerably longer runtimes, suggesting its suitability for exhaustive searches in high-performance computing environments rather than routine screening. Combining all databases (RdRpCATCH-full mode) further increased sensitivity, enabling the broadest possible detection of RdRp sequences; however, this approach also substantially increased the number of FPs-to approximately one in every 200 sequences-which may not be desirable in all contexts.

Furthermore, our findings highlight that bit score thresholds optimized for a particular dataset do not always generalize effectively to more diverse or fragmented samples, such as those commonly found in environmental metatranscriptomic studies. As a result, filtering strategies -including the choice of E-value and bit score cutoffs-should be carefully tailored to the characteristics and requirements of each dataset. In practice, an E-value threshold of 10⁻⁵ offers a robust compromise for general RdRp detection tasks. Under these conditions, the top-performing databases maintained false positive rates below 1 in 2,000 sequences, and sensitivity remained above 0.97 in both tested settings. However, in exploratory analyses focused on discovering novel diversity, relaxing the threshold and using additional validation methods may help identify a wider range of RdRp sequences. Conversely, when minimizing false positives is important, stricter thresholds are recommended (E <10^−9^). In summary, while an E-value threshold of 10⁻⁵ offers a practical and robust compromise for general RdRp detection tasks, the optimal database choice and parameter settings (as well as filtering options based on length or bitscore) will ultimately depend on the specific goals, dataset complexity, and computational constraints of each analysis.

### RdRpCATCH captures novel RdRp diversity in a public metatranscriptomic dataset

To assess RdRpCATCH performance on a publicly available dataset, we reanalyzed a recent study on *Sphagnum* moss viral diversity (46) (see Materials & Methods: ‘Performance assessment versus publicly available datasets’). Building on our results from the comparative analysis scenarios, we evaluated RdRpCATCH using three separate configurations, using different combinations of databases: RdRpCATCH-efficient (RdRp-Scan and Olendraite_fam), RdRpCATCH-balanced (RVMT, Lucaprot_HMM and Zayed_HMM), and RdRpCATCH-full (all the supported databases). For comparison, we deployed geNomad v1.12 to the same dataset, using both conservative and default presets. RdRpCATCH-full, -balanced, and -efficient retrieved 2,596, 2,316, and 1,888 contigs, respectively, while geNomad-default and -conservative, retrieved 6,648 and 1,861(Fig. 6A). Of these, RdRpCATCH-full, -balanced, and -efficient identified 2,033, 1,944, and 1,743 contigs with RdRp evidence or *Orthornavirae* taxonomic annotation, while geNomad-default and -conservative detected 1,880 and 1,553. As expected, most contigs unique to geNomad lacked RdRp evidence, in line with the design of the tools: RdRpCATCH detects RdRp-containing sequences, whereas geNomad captures additional viral sequences without RdRp (e.g., non-RdRp segments, DNA virus transcripts). Notably, most geNomad-default contigs received a ‘Cellular’ annotation, largely reduced in geNomad-conservative, likely due to complex co-assembly composition and classifier limitations. RdRpCATCH found 12, 24, and 105 ‘Cellular’ contigs in efficient, balanced, and full modes, respectively. These findings align with comparative scenarios 1 and 2, as broader database usage enhances sensitivity but also increases false positives. RdRpCATCH-detected cellular-annotated sequences generally had lower bitscores, coverage, and higher E-values (Fig. 6B-D), indicating that stricter thresholds could eliminate most false positives, though some true positives may be lost. Of the RdRpCATCH-full hits, 291 sequences were unclassified by all methods. Their bitscore, E-value, and coverage distributions resemble RdRp-validated sequences (Fig. 6B-D), implying a mixture of diverse RNA viruses and cellular sequences. Further analysis, such as structural similarity searches, could clarify their classification and potentially reveal novel viral diversity.

**Figure 6.**
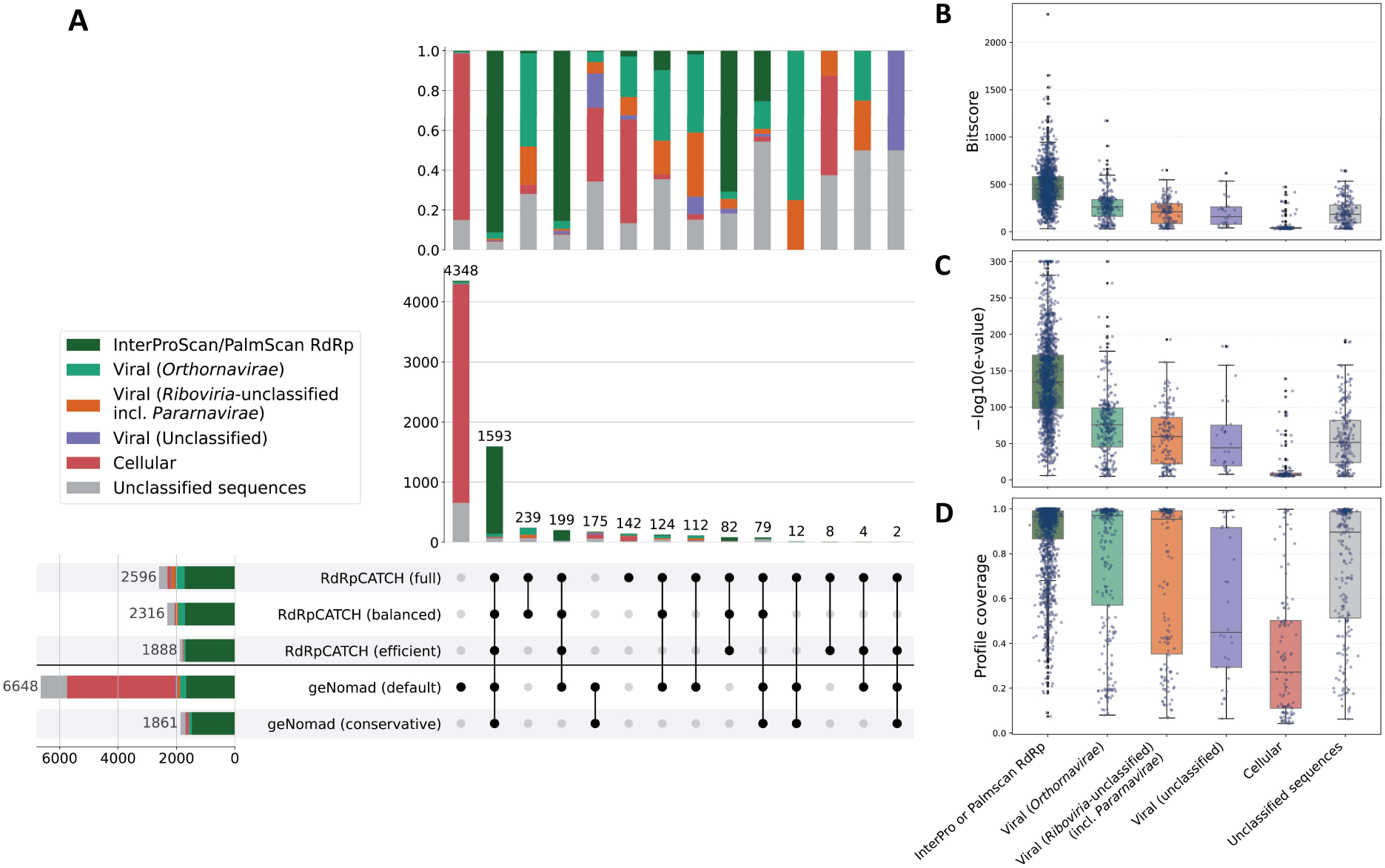
Comparative performance of RdRpCATCH and geNomad and taxonomic annotation per MMSeqs2 easy-taxonomy against Uniref100. (A) Upset plot shows contig-level overlap among RdRpCATCH-full, -efficient, and -balanced configurations and geNomad viral contigs. RdRpCATCH best-hit metrics (B) bitscore, (C) −log 10 e-value, and (D) profile coverage.

## Discussion

The discovery and characterization of RNA viruses in complex metatranscriptomic datasets require robust, sensitive, and scalable detection methods. Most large-scale initiatives aimed at scanning (meta)transcriptomic datasets for RNA viruses rely on the development and application of specialized RNA-dependent RNA polymerase (RdRp) profile hidden Markov model (pHMM) databases. The parallel development of multiple such databases, each designed with distinct principles and optimization strategies, has introduced new challenges related to accessibility, integration into analytical workflows, and the ability to assess performance in controlled environments objectively. RdRpCATCH directly addresses these challenges by offering a unified, user-friendly platform that enables researchers to access and employ multiple publicly available RdRp pHMM resources within a single interface. Furthermore, by providing comprehensive comparative analyses across both canonical and highly diverse datasets, RdRpCATCH delivers insights into the relative strengths and limitations of each approach.

A major feature of RdRpCATCH is its ability to deploy multiple RdRp pHMM databases within a unified framework. This consolidation substantially reduces technical barriers, empowering users to explore datasets for RNA virus signatures efficiently. By automating search, filtering, and annotation, RdRpCATCH enhances reproducibility and transparency in genome-mining studies. The platform supports a range of genetic codes for contig translation and pinpoints the approximate coordinates of the RdRp core domain based on pHMM search results. It prioritizes outputting the sequence-profile pair with the highest bitscore across selected databases and provides provisional taxonomic annotations for positive sequences, facilitating the interpretation of results. Its modular design enables users to incorporate custom databases and seamlessly consolidate their results with those from supported resources. We are committed to maintaining RdRpCATCH as a current and comprehensive resource by integrating new pHMM databases as they become available.

Our results reveal that the choice of database and parameter thresholds should be guided by both the analysis’s goals and complexity, and therefore, we suggest different database sets (RdRpCATCH-balanced, RdRpCATCH-efficient, and RdRpCATCH-full). RdRpCATCH-balanced (Lucaprot_HMM, RVMT, and Zayed_HMM) is well-suited for comprehensive viral discovery across diverse metatranscriptomic datasets when appropriate search parameters are employed. RdRpCATCH-efficient (RdRp-scan and Olendraite_fam) is ideal for high-throughput screening where specificity is paramount. RdRpCATCH-full (all databases) maximizes sensitivity, but with the inevitable increase of FPs. Detailed investigation into the origin of the false positives revealed that profiles (especially from Olendraite_gen, NeoRdRp, and NeoRdRp 2.1) may contain non-RdRp domains (Fig. 4). We considered non-RdRp RNA virus hits to be false positives (FPs) because the interrogated databases were designed exclusively for RdRp detection. However, if these databases are deployed for broader RNA virus discovery, a lower FPR is expected for NeoRdRp, NeoRdRp 2.1, and Olendraite_gen. Users are therefore encouraged to balance sensitivity and specificity according to their research context and to validate key findings through additional annotation methods, phylogenetic analyses, or structural comparison approaches that are well-suited for diverse hits.

The two comparative analysis scenarios and the application of RdRpCATCH to a public dataset demonstrated the tool’s capacity to detect both known and novel viral diversity while maintaining low false positive rates. In a publicly available *Sphagnum* co-assembly, RdRpCATCH identified more sequences with supporting RdRp evidence from additional validation methods or *Orthornavirae* taxonomic annotation compared to geNomad. Furthermore, some sequences detected by RdRpCATCH could not be classified by other methods, suggesting the potential discovery of novel viral diversity. The origins of these unclassified sequences require further clarification through downstream approaches such as structure-based analyses, which have recently been explored to uncover deep evolutionary relationships among RdRps (51). Notably, geNomad incorporates both non-RdRp and RdRp profiles, and utilizes MMSeqs2 to facilitate profile-to-sequence searches. The inclusion of non-RdRp profiles in geNomad may allow for the detection of additional segments not identified by RdRpCATCH, while its RdRp profiles are derived from a range of public databases and several key publications (e.g., RVMT). Since RVMT is integrated into both geNomad and RdRpCATCH, this likely accounts for the observed overlap in their results (Fig. 6A). The extra RdRp-containing hits of RdRpCATCH, however, can be attributed to its use of a broader array of RdRp databases, the increased sensitivity of pHMMs compared to MMSeqs2, and differences in translation strategies (with RdRpCATCH employing six-frame translation, whereas geNomad uses prodigal-gv).

While the comparative analysis scenarios provide valuable insights for the performance of RdRpCATCH and the underlying databases, we acknowledge that methodological limitations can impact interpretation of the results. First, the RdRp sequences used for the comparative analysis scenario 2 are not validated or accepted by ICTV; rather, they have been described as RdRps in previous studies, and our goal was to evaluate the pHMM databases’ capacity to detect them. Additionally, many of the sequences included in this dataset were used as seed sequences for constructing the pHMMs, and thus generalization to novel taxa that are not represented in the seed sequences is expected to be less accurate or reliable. Likewise, the negative datasets selected for our analysis were intended to represent a broad range of sequences potentially present in RNA-Seq assemblies. Nevertheless, complex and diverse assemblies may harbour various cellular-origin sequences absent from these datasets, potentially resulting in more false positives than anticipated. Notably, RdRp-based methods can detect both *bona fide* exogenous RNA viruses and non-retroviral endogenous viral elements (nr-EVEs), as highlighted in recent systematic analyses of mosquito transcriptomes and public repositories (52, 53). Although not assessed in this manuscript, it is expected that the supported databases can yield nr-EVEs, which may inflate apparent virus richness and downstream ecological interpretations. If distinguishing between nr-EVEs and exogenous RNA viruses is necessary, we recommend assessing the similarity of RdRp-candidate contigs to host genome reference assemblies or employing specialized tools for endogenous viral element detection (54, 55).

As part of RdRpCATCH, we provide provisional taxonomic annotations for predicted viral sequences. Our annotation method is efficient and based on similarity searches against a well-defined NCBI RefSeq database of RNA viral proteins. However, this method is not exhaustive and may not yield annotations for novel lineages, particularly for diverse RNA virus sequences not included in the RefSeq database. Therefore, we recommend that taxonomic annotations for these sequences are validated downstream, either manually based on phylogenetic analysis or by employing specialized automated frameworks designed for the taxonomic annotation of metagenomically detected RNA virus sequences (43, 56, 57). For post-processing sequences detected by pHMM databases, most model developers validate the presence of three core catalytic motifs of the palm subdomain in the yielded sequences, either through manual review or semi-automatically using multiple sequence alignments (16, 23, 25). These motifs (A, B, and C) are conserved in most RdRps and are crucial for catalyzing the phosphodiester bond formation during RNA chain elongation and for template binding (58). As a general guideline, a high profile coverage may indicate the presence of catalytic motifs in the sequence. The PalmScan tool can automatically detect the A, B, and C motifs and their configurations (21); however, it is optimized for precision and may occasionally lack the sensitivity required to identify motifs in divergent RdRps, as showcased for divergent ormycoviruses (59). Therefore, sequences not identified as positive by PalmScan should undergo a second round of screening, with a manual review and/or structural comparison to confirm the presence of core motifs.

RdRpCATCH focuses on detecting the RdRp sequence in metatranscriptomic datasets to facilitate the discovery of RNA viruses, and its profiles are highly specialized for this task. Notably, our taxonomic analysis of false positive assignments suggests that some profile models may inadvertently capture non-RdRp regions or unclassified viral sequences, highlighting an area for future refinement in database curation. Some resources focus on RNA virus detection by integrating more RNA virus protein families, such as VirBot (60), or more generalistic profile databases that can be used for both DNA and RNA virus discovery tasks (61). While these approaches offer the advantage of capturing a wider array of viral signatures, integrating them into a unified detection framework may require further optimization to balance sensitivity, specificity, and computational efficiency.

A natural limitation of RdRp-based virus discovery is that only RdRp-containing sequences can be found, whereas other segments of segmented and multipartite viruses cannot be detected. This approach provides an advantage for ecological studies, where the restriction to one segment prevents the repeated inclusion of multiple segments belonging to the same genome when assessing virus diversity. However, detecting only the RdRp-containing segment does not provide the full genomic context of segmented and multipartite viruses. To this end, emerging methods are based on common co-occurrence and similar abundances of segments (62, 63), on taxonomic relationships of segments with known fragmented viral genomes (64), or on detecting the conserved 5’ and 3’ UTRs and mining them from a given metatranscriptomics dataset (65). Furthermore, conventional similarity based-methods (i.e. BLAST, MMSeqs) and profile-based methods, such as the ones discussed in the previous paragraph, can be effective in retrieving non-RdRp segments. In such cases, the manual assignment of segments to the same genome can be challenging, as it requires knowledge of the genome architecture of the corresponding viruses and it potentially excludes novel segments that do not show similarity to existing sequences in the target databases. All of the above are viable options for this task; however, these approaches are each virus taxon specific and this task has not yet been conclusively addressed, especially for novel viruses whose genome architecture remains unverified.

Deep learning approaches have demonstrated considerable potential in advancing the discovery of RNA viruses from metatranscriptomic data. Recently, several models with various architectures have been developed. Some of these models are designed for specific purposes; for instance, VirDetect-AI is focused on eukaryotic viruses, while VirHunter targets plant transcriptomes (18, 66). Other models, such as ViralLM, have a broader scope, enabling detection of both DNA and RNA viruses (17). Notably, Lucaprot specifically emphasizes RdRp detection and employs a transformer-based architecture to integrate both sequence and structural information (16). While these deep learning methods offer promising avenues for RNA virus discovery, profile Hidden Markov Models (pHMMs) have demonstrated exceptional efficiency and high performance in capturing divergent RNA virus signals, and novel methods should be benchmarked against established pHMM approaches to assess their performance. Additionally, the interpretability and biological relevance of pHMM results provide valuable insights into viral relationships and evolutionary patterns and training of ML models is much more computationally expensive than construction of pHMMs. This makes them particularly relevant for comprehensive RNA virus discovery workflows, where accuracy, speed, and biological interpretability are essential considerations.

In conclusion, RdRpCATCH offers a unified framework for RdRp-based RNA virus discovery, substantially reducing technical barriers and facilitating a systematic exploration of the global RNA virosphere. Our comparative analysis offers practical guidance for selecting databases and parameters, emphasizing the importance of a tailored approach informed by dataset characteristics and research objectives. Continued development and community-driven standardization will further enhance the utility and impact of genome mining strategies in RNA virology.

## Data availability

The databases that are used in this study are available in Zenodo (https://doi.org/10.5281/zenodo.15463729). The code for RdRpCATCH is publicly available on GitHub (https://github.com/dimitris-karapliafis/RdRpCATCH) and is archived in Zenodo (https://doi.org/10.5281/zenodo.20613411). The code for RdRpCATCH is publicly available in GitHub (https://github.com/dimitris-karapliafis/RdRpCATCH). The web server is available at https://rdrpcatch.bioinformatics.nl/. The comparative analysis datasets, code, and results are stored in Zenodo (https://doi.org/10.5281/zenodo.18470715)

## Author contributions

D.K.: Conceptualization, Methodology, Investigation, Formal Analysis, Data Curation, Software, Visualization, Writing - Original Draft, Writing - Review & Editing

U.N.: Conceptualization, Data Curation, Software, Writing – Review & Editing

I.O.: Conceptualization, Data Curation, Writing – Review & Editing

J.C.: Conceptualization, Writing – Review & Editing

S.S.: Conceptualization, Writing – Review & Editing

X.H.: Writing – Review & Editing

D.d.R.: Writing – Review & Editing, Resources, Supervision

M.Z.: Writing – Review & Editing, Supervision

A.K: Conceptualization, Writing – Review & Editing, Resources, Supervision, Project Administration, Funding Acquisition

## Acknowledgments

We thank Marco Forgia, Hisham Shaikh and Ella Sieradzki for their valuable feedback during the design phase, as well as the broader RdRp Summit community for their ongoing support. We also thank Astrid Bryon, Mattias de Hollander and Laura Patiño Medina for testing the RdRpCATCH package, Arjan Draisma for assistance with web server deployment, and Dirk-Jan van Workum for expert advice on Bioconda packaging.

This work was supported by the Graduate School Experimental Plant Sciences (EPS), the Netherlands [3184519001]. The work conducted by the U.S. Department of Energy Joint Genome Institute (https://ror.org/04xm1d337), a DOE Office of Science User Facility, is supported by the Office of Science of the U.S. Department of Energy operated under Contract No. DE-AC02-05CH11231.

## Notes

https://github.com/dimitris-karapliafis/RdRpCATCH

https://doi.org/10.5281/zenodo.18470715

https://doi.org/10.5281/zenodo.15463729

https://doi.org/10.5281/zenodo.20613411

### Competing Interest Statement

The authors have declared no competing interest.

### Summary of Updates

We added an additional analysis part that assesses the performance of RdRpCATCH when deployed on a publicly available dataset.

## References

1. Hendrix, R.W., Smith, M.C.M., Burns, R.N., Ford, M.E. and Hatfull, G.F. (1999) Evolutionary relationships among diverse bacteriophages and prophages: All the world’s a phage. Proc. Natl. Acad. Sci., 96, 2192–2197.

2. Mushegian, A.R. (2020) Are There 1031 Virus Particles on Earth, or More, or Fewer? J. Bacteriol., 202, e00052–20.

3. Koonin, E.V., Krupovic, M. and Agol, V.I. (2021) The Baltimore Classification of Viruses 50 Years Later: How Does It Stand in the Light of Virus Evolution? Microbiol. Mol. Biol. Rev. MMBR, 85, e00053–21.

4. Duffy, S. (2018) Why are RNA virus mutation rates so damn high? PLoS Biol., 16, e3000003.

5. Payne, S. (2017) Introduction to RNA Viruses. Viruses, 10.1016/B978-0-12-803109-4.00010-6.

6. Elena, S.F. and Sanjuán, R. (2005) Adaptive Value of High Mutation Rates of RNA Viruses: Separating Causes from Consequences. J. Virol., 79, 11555–11558.

7. Simon-Loriere, E. and Holmes, E.C. (2011) Why do RNA viruses recombine? Nat. Rev. Microbiol., 9, 617–626.

8. Black, E.J., Powell, C.S., Dempsey, D.M., Hendrickson, R.C., Mims, L.R. and Lefkowitz, E.J. (2025) Virus taxonomy: the database of the International Committee on Taxonomy of Viruses. Nucleic Acids Res., 10.1093/nar/gkaf1159.

9. Nakagawa, S., Sakaguchi, S., Ogura, A., Mineta, K., Endo, T., Suzuki, Y. and Gojobori, T. (2023) Current trends in RNA virus detection through metatranscriptome sequencing data. FEBS Open Bio, 13, 992–1000.

10. Simmonds, P., Adams, M.J., Benkő, M., Breitbart, M., Brister, J.R., Carstens, E.B., Davison, A.J., Delwart, E., Gorbalenya, A.E., Harrach, B., et al. (2017) Virus taxonomy in the age of metagenomics. Nat. Rev. Microbiol., 15, 161–168.

11. Dominguez-Huerta, G., Wainaina, J.M., Zayed, A.A., Culley, A.I., Kuhn, J.H. and Sullivan, M.B. (2023) The RNA virosphere: How big and diverse is it? Environ. Microbiol., 25, 209.

12. Koonin, E.V., Krupovic, M. and Dolja, V.V. (2023) The global virome: How much diversity and how many independent origins? Environ. Microbiol., 25, 40–44.

13. Mönttinen, H.A.M., Ravantti, J.J. and Poranen, M.M. (2021) Structure Unveils Relationships between RNA Virus Polymerases. Viruses, 13, 313.

14. Bruenn, J.A. (2003) A structural and primary sequence comparison of the viral RNA-dependent RNA polymerases. Nucleic Acids Res., 31, 1821–1829.

15. Skewes-Cox, P., Sharpton, T.J., Pollard, K.S. and DeRisi, J.L. (2014) Profile Hidden Markov Models for the Detection of Viruses within Metagenomic Sequence Data. PLOS ONE, 9, e105067.

16. Hou, X., He, Y., Fang, P., Mei, S.-Q., Xu, Z., Wu, W.-C., Tian, J.-H., Zhang, S., Zeng, Z.-Y., Gou,Q.-Y., et al. (2024) Using artificial intelligence to document the hidden RNA virosphere. Cell, 187, 6929–6942.e16.

17. Peng, C., Shang, J., Guan, J., Wang, D. and Sun, Y. (2024) ViraLM: empowering virus discovery through the genome foundation model. Bioinformatics, 40, btae704.

18. Zárate, A., Díaz-González, L. and Taboada, B. (2025) VirDetect-AI: a residual and convolutional neural network–based metagenomic tool for eukaryotic viral protein identification. Brief. Bioinform., 26, bbaf001.

19. Park, J., Karplus, K., Barrett, C., Hughey, R., Haussler, D., Hubbard,T. and Chothia,C. (1998) Sequence comparisons using multiple sequences detect three times as many remote homologues as pairwise methods1. J. Mol. Biol., 284, 1201–1210.

20. Wolf, Y.I., Silas, S., Wang, Y., Wu, S., Bocek, M., Kazlauskas, D., Krupovic, M., Fire, A., Dolja, V.V. and Koonin, E.V. (2020) Doubling of the known set of RNA viruses by metagenomic analysis of an aquatic virome. Nat. Microbiol., 5, 1262–1270.

21. Babaian, A. and Edgar, R. (2022) Ribovirus classification by a polymerase barcode sequence. PeerJ, 10, e14055.

22. Edgar, R.C., Taylor, B., Lin, V., Altman, T., Barbera, P., Meleshko, D., Lohr, D., Novakovsky, G., Buchfink, B., Al-Shayeb, B., et al. (2022) Petabase-scale sequence alignment catalyses viral discovery. Nature, 602, 142–147.

23. Neri, U., Wolf, Y.I., Roux, S., Camargo, A.P., Lee, B., Kazlauskas, D., Chen, I.M., Ivanova, N., Allen, L.Z., Paez-Espino, D., et al. (2022) Expansion of the global RNA virome reveals diverse clades of bacteriophages. Cell, 185, 4023–4037.e18.

24. Zayed, A.A., Wainaina, J.M., Dominguez-Huerta, G., Pelletier, E., Guo, J., Mohssen, M., Tian, F., Pratama, A.A., Bolduc, B., Zablocki, O., et al. (2022) Cryptic and abundant marine viruses at the evolutionary origins of Earth’s RNA virome. Science, 376, 156–162.

25. Olendraite, I., Brown, K. and Firth, A.E. (2023) Identification of RNA Virus–Derived RdRp Sequences in Publicly Available Transcriptomic Data Sets. Mol. Biol. Evol., 40, msad060.

26. Charon, J., Buchmann, J.P., Sadiq, S. and Holmes, E.C. (2022) RdRp-scan: A bioinformatic resource to identify and annotate divergent RNA viruses in metagenomic sequence data. Virus Evol., 8, veac082.

27. Sakaguchi, S., Nakano, T. and Nakagawa, S. (2024) NeoRdRp2 with improved seed data, annotations, and scoring. Front. Virol., 4.

28. Charon, J., Olendraite, I., Forgia, M., Chong, L.C., Hillary, L.S., Roux, S., Kupczok, A., Debat, H., Sakaguchi, S., Tahzima, R., et al. (2024) Consensus statement from the first RdRp Summit: advancing RNA virus discovery at scale across communities. Front. Virol., 4.

29. Sakaguchi, S., Urayama, S., Takaki, Y., Hirosuna, K., Wu, H., Suzuki, Y., Nunoura, T., Nakano, T. and Nakagawa, S. (2022) NeoRdRp: A Comprehensive Dataset for Identifying RNA-dependent RNA Polymerases of Various RNA Viruses from Metatranscriptomic Data. Microbes Environ., 37, ME22001.

30. Aiewsakun, P. and Simmonds, P. (2018) The genomic underpinnings of eukaryotic virus taxonomy: creating a sequence-based framework for family-level virus classification. Microbiome, 6, 38.

31. Olendraite, I. (2021) Mining Diverse and Novel RNA Viruses in Transcriptomic Datasets. 10.17863/CAM.76470.

32. Eddy, S.R. (2011) Accelerated Profile HMM Searches. PLOS Comput. Biol., 7, e1002195.

33. Paysan-Lafosse, T., Andreeva, A., Blum, M., Chuguransky, S.R., Grego, T., Pinto, B.L., Salazar, G.A., Bileschi, M.L., Llinares-López, F., Meng-Papaxanthos,L., et al. (2025) The Pfam protein families database: embracing AI/ML. Nucleic Acids Res., 53, D523–D534.

34. Tian, Z., Hu, T., Holmes, E.C., Ji, J. and Shi, W. (2024) Analysis of the genetic diversity in RNA-directed RNA polymerase sequences: implications for an automated RNA virus classification system. Virus Evol., 10, veae059.

35. Brown, K. and Firth, A.E. (2025) Uncovering hundreds of exogenous and endogenous RNA viral RdRp sequences amongst uncharacterized sequences in public protein databases. Virus Evol., 11, veaf074.

36. Blum, M., Andreeva, A., Florentino, L.C., Chuguransky, S.R., Grego, T., Hobbs, E., Pinto, B.L., Orr, A., Paysan-Lafosse, T., Ponamareva, I., et al. (2025) InterPro: the protein sequence classification resource in 2025. Nucleic Acids Res., 53, D444–D456.

37. Larralde, M. and Zeller, G. (2023) PyHMMER: a Python library binding to HMMER for efficient sequence analysis. Bioinformatics, 39, btad214.

38. VanderPlas, J., Granger, B.E., Heer, J., Moritz, D., Wongsuphasawat, K., Satyanarayan, A., Lees, E., Timofeev, I., Welsh, B. and Sievert, S. (2018) Altair: Interactive Statistical Visualizations for Python. J. Open Source Softw., 3, 1057.

39. Shen, W., Le, S., Li, Y. and Hu, F. (2016) SeqKit: A Cross-Platform and Ultrafast Toolkit for FASTA/Q File Manipulation. PLOS ONE, 11, e0163962.

40. Mirdita, M., Steinegger, M., Breitwieser, F., Söding, J. and Levy Karin, E. (2021) Fast and sensitive taxonomic assignment to metagenomic contigs. Bioinformatics, 37, 3029–3031.

41. Steinegger, M. and Söding, J. (2017) MMseqs2 enables sensitive protein sequence searching for the analysis of massive data sets. Nat. Biotechnol., 35, 1026–1028.

42. Hingamp, P., Grimsley, N., Acinas, S.G., Clerissi, C., Subirana, L., Poulain, J., Ferrera, I., Sarmento, H., Villar, E., Lima-Mendez, G., et al. (2013) Exploring nucleo-cytoplasmic large DNA viruses in Tara Oceans microbial metagenomes. ISME J., 7, 1678–1695.

43. Yang, Z., Shan, Y., Liu, X., Chen, G., Pan, Y., Gou, Q., Zou, J., Chang, Z., Zeng, Q., Yang, C., et al. (2024) VirID: Beyond Virus Discovery—An Integrated Platform for Comprehensive RNA Virus Characterization. Mol. Biol. Evol., 41, msae202.

44. The UniProt Consortium (2025) UniProt: the Universal Protein Knowledgebase in 2025. Nucleic Acids Res., 53, D609–D617.

45. Ahmad, S., Jose da Costa Gonzales, L., Bowler-Barnett, E.H., Rice, D.L., Kim, M., Wijerathne, S., Luciani, A., Kandasaamy, S., Luo, J., Watkins, X., et al. (2025) The UniProt website API: facilitating programmatic access to protein knowledge. Nucleic Acids Res., 53, W547–W553.

46. Denison, E.R., Pound, H.L., Gann, E.R., Gilbert, N.E., Weston, D.J., Pelletier, D.A. and Wilhelm, S.W. (2025) Identification of shared viral sequences in peat moss metagenomes reveals elements of a possible Sphagnum core virome. Environ. Microbiome, 20, 62.

47. Li, D., Liu, C.-M., Luo, R., Sadakane, K. and Lam, T.-W. (2015) MEGAHIT: an ultra-fast single-node solution for large and complex metagenomics assembly via succinct de Bruijn graph. Bioinformatics, 31, 1674–1676.

48. Camargo, A.P., Roux, S., Schulz, F., Babinski, M., Xu, Y., Hu, B., Chain, P.S.G., Nayfach, S. and Kyrpides, N.C. (2023) Identification of mobile genetic elements with geNomad. Nat. Biotechnol., 10.1038/s41587-023-01953-y.

49. Jones, P., Binns, D., Chang, H.-Y., Fraser, M., Li, W., McAnulla, C., McWilliam, H., Maslen, J., Mitchell, A., Nuka, G., et al. (2014) InterProScan 5: genome-scale protein function classification. Bioinformatics, 30, 1236–1240.

50. Nibert, M.L. (2017) Mitovirus UGA(Trp) codon usage parallels that of host mitochondria. Virology, 507, 96–100.

51. Mönttinen, H.A.M., Ravantti, J.J., Mayne, R., Simmonds, P. and Poranen, M.M. (2026) Revealing deep evolutionary relationships between RNA viruses using predicted structural models of viral RNA polymerases. Mol. Biol. Evol., 43, msag088.

52. Brown, K. and Firth, A.E. (2025) Uncovering hundreds of exogenous and endogenous RNA viral RdRp sequences amongst uncharacterized sequences in public protein databases. Virus Evol., 11, veaf074.

53. Brait, N., Hackl, T., Morel, C., Exbrayat, A., Gutierrez, S. and Lequime, S. (2024) A tale of caution: How endogenous viral elements affect virus discovery in transcriptomic data. Virus Evol., 10, vead088.

54. Brait, N., Hackl, T. and Lequime, S. (2025) detectEVE: Fast, Sensitive and Precise Detection of Endogenous Viral Elements in Genomic Data. Mol. Ecol. Resour., 25, e14083.

55. Muñoz-Baena, L., Harding, E.F., Nino Barreat, J.G., Kinsella, C.M. and Katzourakis, A. (2025) HI-FEVER: a Nextflow pipeline for the high-throughput discovery and annotation of endogenous viral elements. Bioinformatics, 41, btaf610.

56. Zheng, K., Sun, J., Liang, Y., Kong, L., Paez-Espino, D., Mcminn, A. and Wang, M. (2025) VITAP: a high precision tool for DNA and RNA viral classification based on meta-omic data. Nat. Commun., 16, 2226.

57. Zhu, Y., Chen, G. and Sun, Y. (2024) VirTAXA: enhancing RNA virus taxonomic classification with remote homology search and tree-based validation. Bioinformatics, 40, btae575.

58. te Velthuis, A.J.W. (2014) Common and unique features of viral RNA-dependent polymerases. Cell. Mol. Life Sci. CMLS, 71, 4403–4420.

59. Forgia, M., Chiapello, M., Daghino, S., Pacifico, D., Crucitti, D., Oliva, D., Ayllon, M., Turina, M. and Turina, M. (2022) Three new clades of putative viral RNA-dependent RNA polymerases with rare or unique catalytic triads discovered in libraries of ORFans from powdery mildews and the yeast of oenological interest Starmerella bacillaris. Virus Evol., 8, veac038.

60. Chen, G., Tang, X., Shi, M. and Sun, Y. (2023) VirBot: an RNA viral contig detector for metagenomic data. Bioinformatics, 39, btad093.

61. Yu, R., Huang, Z., Lam, T.Y.C. and Sun, Y. (2024) Utilizing profile hidden Markov model databases for discovering viruses from metagenomic data: a comprehensive review. Brief. Bioinform., 25, bbae292.

62. Liu, X., Kong, J., Shan, Y., Yang, Z., Miao, J., Pan, Y., Luo, T., Shi, Z., Wang, Y., Gou, Q., et al. (2025) SegFinder: an automated tool for identifying complete RNA virus genome segments through co-occurrence in multiple sequenced samples. Brief. Bioinform., 26, bbaf358.

63. Obbard, D.J., Shi, M., Roberts, K.E., Longdon, B. and Dennis, A.B. (2020) A new lineage of segmented RNA viruses infecting animals. Virus Evol., 6, vez061.

64. Tang, X., Shang, J., Chen, G., Chan, K.H.K., Shi, M. and Sun, Y. (2024) SegVir: Reconstruction of Complete Segmented RNA Viral Genomes from Metatranscriptomes. Mol. Biol. Evol., 41, msae171.

65. Zhang, S., Yang, C., Qiu, Y., Liao, R., Xuan, Z., Ren, F., Dong, Y., Xie, X., Han, Y., Wu, D., et al. (2024) Conserved untranslated regions of multipartite viruses: Natural markers of novel viral genomic components and tags of viral evolution. Virus Evol., 10, veae004.

66. Sukhorukov, G., Khalili, M., Gascuel, O., Candresse, T., Marais-Colombel, A. and Nikolski, M. (2022) VirHunter: A Deep Learning-Based Method for Detection of Novel RNA Viruses in Plant Sequencing Data. Front. Bioinforma., 2.

