## Supplementary material for "RdRpCATCH: A unified resource for RNA virus discovery using viral RNA-dependent RNA polymerase profile Hidden Markov models"

* - corresponding author

### Supplementary material

#### Table of contents

Table S1. Databases supported by RdRpCATCH and their respective sources from which the pHMM databases were retrieved. ........ 3
Table S2. Summary of the RdRp datasets that were used to compare the performance of the databases, the resources from which the sequences were retrieved, and a brief description of their composition. ........ 4
Table S3. Summary of the negative sequence datasets that were used to compare the performance of the databases, the resources from which the sequences were retrieved, and a brief description of their composition. 5
Figure S1.(First column) Distribution of the number of sequences used to construct each profile-HMM. (Second column) Distribution of the lengths of the profile-HMMs. (Third column) Scatter plot illustrating the relationship between profile-HMM length and the number of sequences used for its construction. ........ 6
Figure S2. Bitscore distributions of true and false positive hits across different E-value thresholds and HMM databases for the comparative analysis scenario 1 (VirID_TP-Lucaprot_TN dataset). Each panel corresponds to a specific HMM database. For each E-value, the optimal bitscore threshold was determined by maximizing the F1-score. An annotation arrow highlights the overall best-performing E-value and its corresponding optimal bitscore, with the text box displaying the maximum F1-score achieved for that database. Dotted orange lines indicate bitscore thresholds for other E-values that also achieved this maximum F1-score (tie-break candidates), while dotted black lines show the optimal bitscores for the remaining, sub-optimal E-value thresholds ........ 7
Figure S6. Bitscore distributions of true and false positive hits across different E-value thresholds and HMM databases for comparative analysis scenario 2 (Total_TP/Total_TN dataset). Each panel corresponds to a specific HMM database. For each E-value, the optimal bitscore threshold was determined by maximizing the F1-score. An annotation arrow highlights the overall best-performing E-value and its corresponding optimal bitscore, with the text box displaying the maximum F1-score achieved for that database. Dotted orange lines indicate bitscore thresholds for other E-values that also achieved this maximum F1-score (tie-break candidates), while dotted black lines show the optimal bitscores for the remaining, sub-optimal E-value thresholds. ........ 11

Table S1. Databases supported by RdRpCATCH and their respective sources from which the pHMM databases were retrieved.

| Name | Number of profiles and source | Source |
|---|---|---|
| **Lucaprot_HMM** | 754 pHMMs generated by iteratively interrogating 10,487 metatranscriptomes (1) | https://figshare.com/articles/thesis/The_LucaProt-Related_Resources/26298802/16 |
| **NeoRdRp** | 1,182 pHMMs generated from 12,502 RdRp domain sequences (2) | https://github.com/shoichisakaguchi/NeoRdRp |
| **NeoRdRp 2.1** | 19,394 high-granularity pHMMs derived from various studies (3) | https://zenodo.org/records/10851672 |
| **Olendraite_fam** | 77 family-level pHMMs identified in the transcriptome shotgun assembly sequence database (TSA) (4) | https://github.com/ingridole/ViralRdRp_pHMMs |
| **Olendraite_gen** | 341 genus-level pHMMs identified in the transcriptome shotgun assembly sequence database (TSA) (5). | https://doi.org/10.17863/CAM.76470 |
| **RdRp-scan** | 68 curated RdRP pHMMs for the detection of divergent RNA viruses (6) | https://github.com/JustineCharon/RdRp-scan |
| **RVMT** | 710 RdRp pHMMs generated by mining viruses in 3,598 diverse metatranscriptomes (7) | https://github.com/UriNeri/ColabScan/tree/main/DBs |
| **Zayed_HMM** | 2,489 pHMMs generated by iteratively interrogating the Tara Oceans datasets (8) | https://datacommons.cyverse.org/browse/iplant/home/shared/iVirus/ZayedWainainaDominguez-Huerta_RNAevolution_Dec2021 |

Table S2. Summary of the RdRp datasets that were used to compare the performance of the databases, the resources from which the sequences were retrieved, and a brief description of their composition.

| Dataset name | Number of sequences | Source | Brief description |
|---|---|---|---|
| Hou_RdRps | 161,979 | https://figshare.com/articles/thesis/The_LucaProt-Related_Resources/26298802 | Curated set of RdRp sequences derived from 10,487 metatranscriptomes (1). |
| Neri_RdRps | 77,510 | https://zenodo.org/records/7368133 | Curated set of full-length RdRP core domain sequences derived from 5,150 metatranscriptomes (7). |
| Olendraite_RdRps | 12,110 | https://github.com/ingridole/ViralRdRp_pHMMs_2 | Curated set of ORFs containing RdRps or RdRp fragments from Transcriptome Shotgun Assembly database (4). |
| PalmDB_RdRps | 513,176 | https://github.com/ababaian/palmdb | Set of RdRp palmprint sequences clustered at 90% amino acid identity (9). |
| VirID_core_RdRps | 7,080 | https://github.com/ZiyueYang01/VirID/tree/main | Curated set of RdRp sequences derived from NCBI Genbank, NCBI RefSeq and backbone RdRp set (10) |
| Wolf_RdRps | 4,593 | https://ftp.ncbi.nih.gov/pub/wolf/_suppl/yangshan/ | Curated set of RdRps derived from Yangshan Deep-Water Harbour metatranscriptome (11) |
| Zayed_RdRps | 6,238 | https://datacommons.cyverse.org/browse/iplant/home/shared/iVirus/ZayedWainainaDominguez-Huerta_RNAevolution_Dec2021 | Curated set of RdRp core sequences derived from Tara Oceans metatranscriptomes (8). |

Table S3. Summary of the negative sequence datasets that were used to compare the performance of the databases, the resources from which the sequences were retrieved, and a brief description of their composition.

| Dataset name | Number of sequences | Source | Brief description |
|---|---|---|---|
| Lucaprot_TN | 229,434 | https://figshare.com/articles/thesis/The_LucaProt-Related_Resources/26298802 | A dataset of non-viral RdRp sequences formulated as part of the Lucaprot study (1). |
| custom_TN | 576,624 | https://www.uniprot.org/ | A wide array of cellular and non-orthornavirae proteins retrieved from SwissProt (12). |

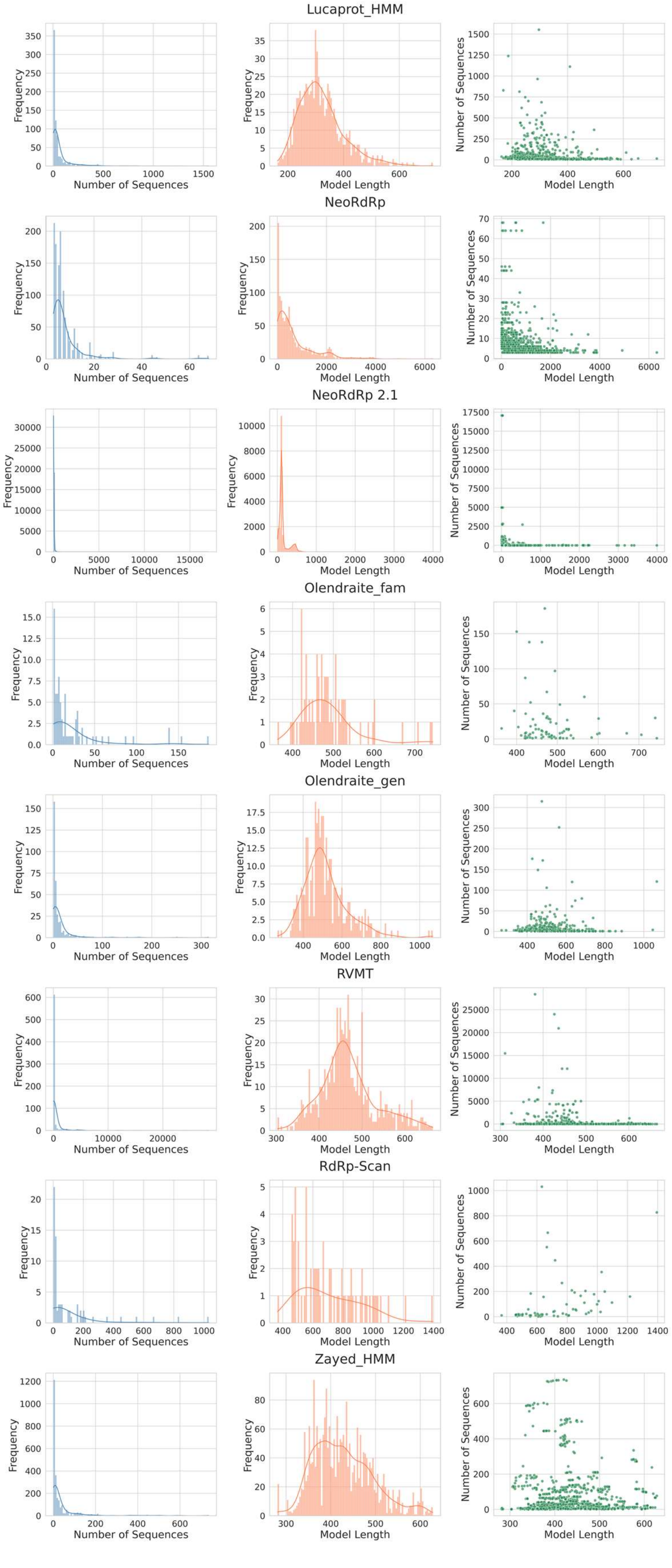

Figure S1. (First column) Distribution of the number of sequences used to construct each profile-HMM. (Second column) Distribution of the lengths of the profile-HMMs. (Third column) Scatter plot illustrating the relationship between profile-HMM length and the number of sequences used for its construction.

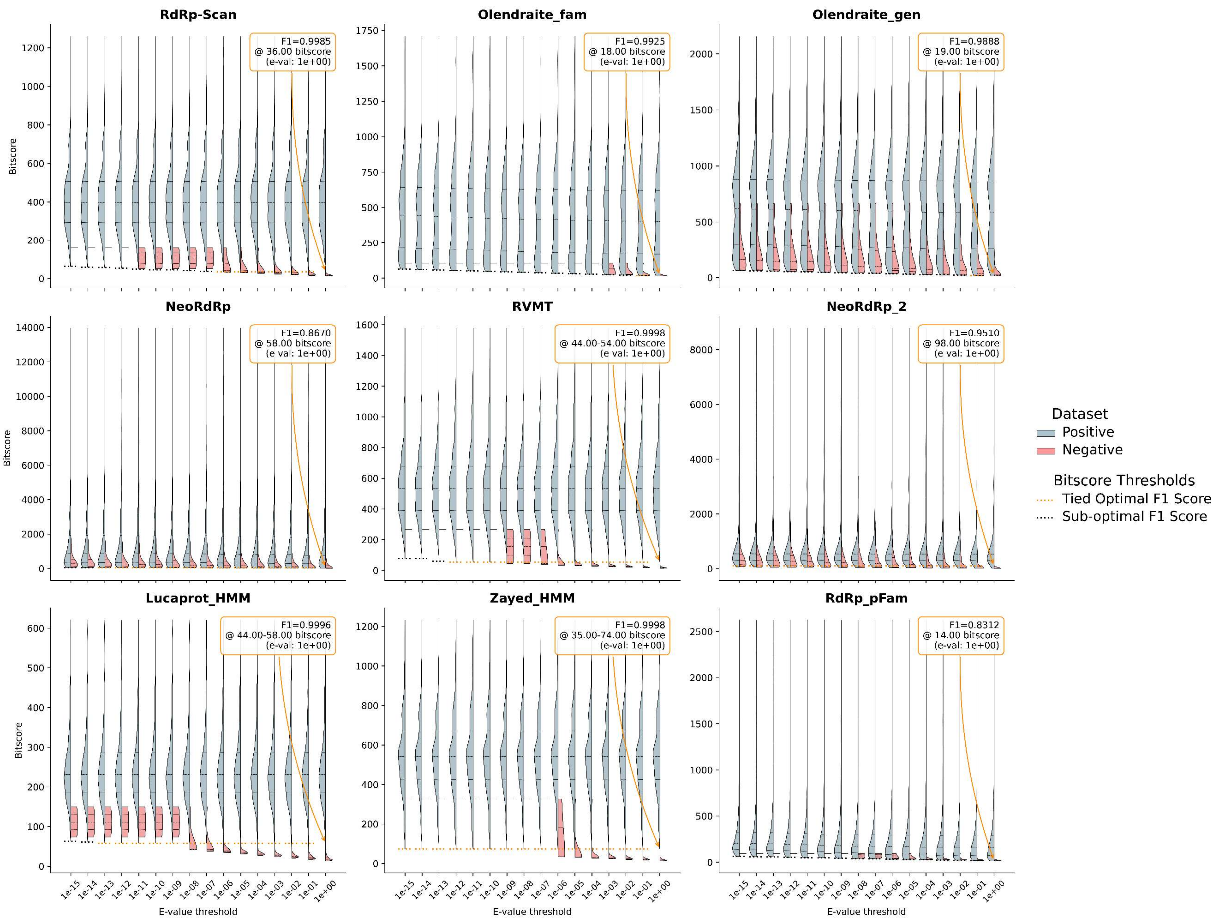

Figure S2. Bitscore distributions of true and false positive hits across different E-value thresholds and HMM databases for the comparative analysis scenario 1 (VirID_TP/Lucaprot_TN datasets). Each panel corresponds to a specific HMM database. For each E-value, the optimal bitscore threshold was determined by maximizing the F1-score. An annotation arrow highlights the overall best-performing E-value per database and its corresponding optimal bitscore, with the text box displaying the maximum F1-score achieved for that database. Dotted orange lines indicate bitscore thresholds for other E-values that also achieved this maximum F1-score (tie-break candidates), while dotted black lines show the optimal bitscores for the remaining, sub-optimal E-value thresholds.

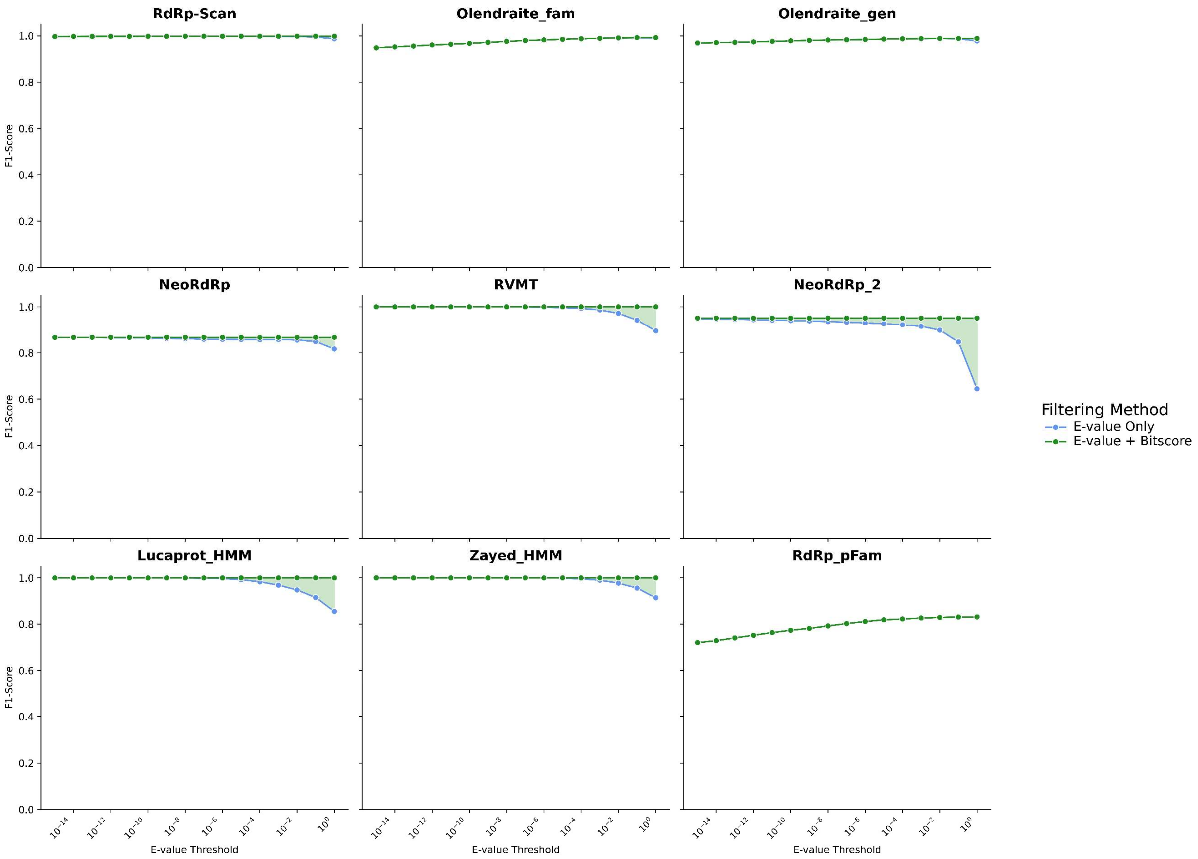

Figure S3. Improvement in F1-score after applying optimized bitscore filtering for comparative analysis scenario 1 (VirID_TP/Lucaprot_TN datasets). Each panel displays the performance for a single HMM database across a range of E-value thresholds (x-axis, log scale).

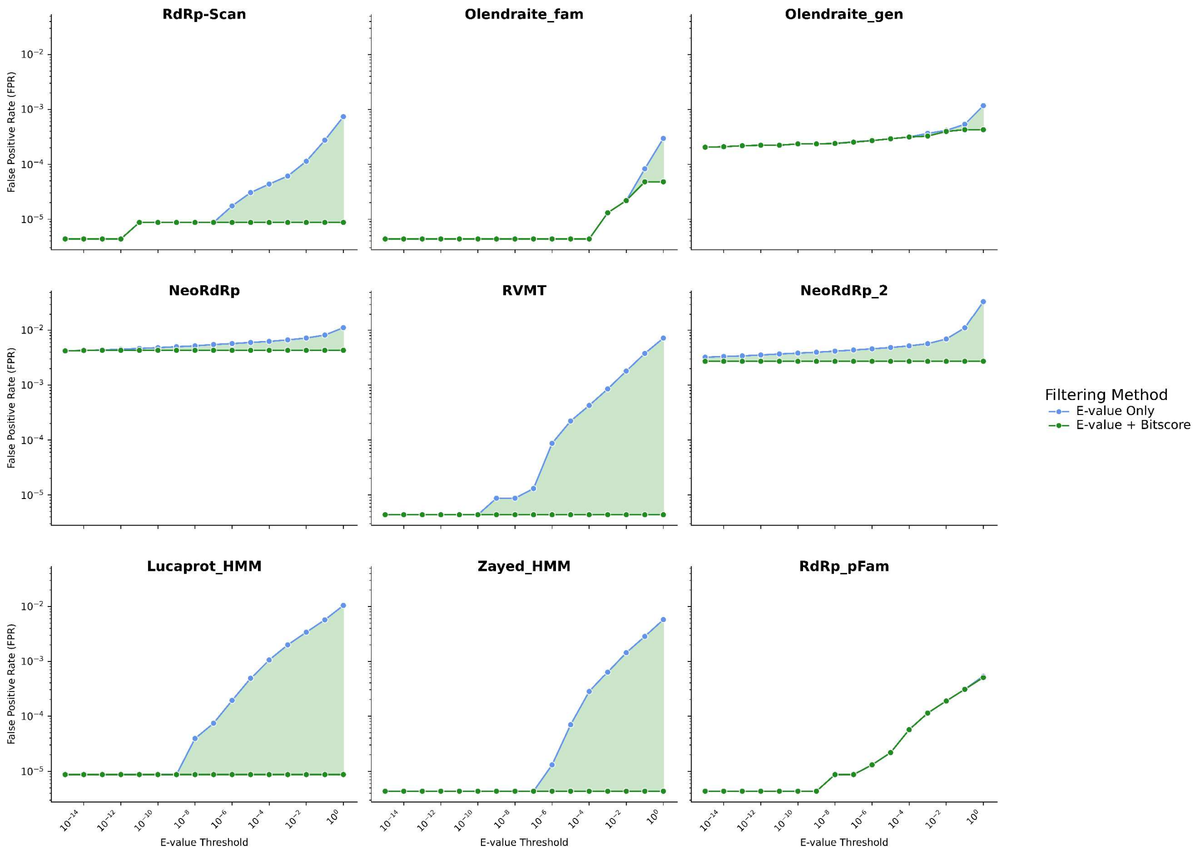

Figure S4. Improvement in FPR after applying optimized bitscore filtering for comparative analysis scenario 1 (VirID_TP/Lucaprot_TN datasets). Each panel displays the performance for a single HMM database across a range of E-value thresholds (x-axis, log scale).

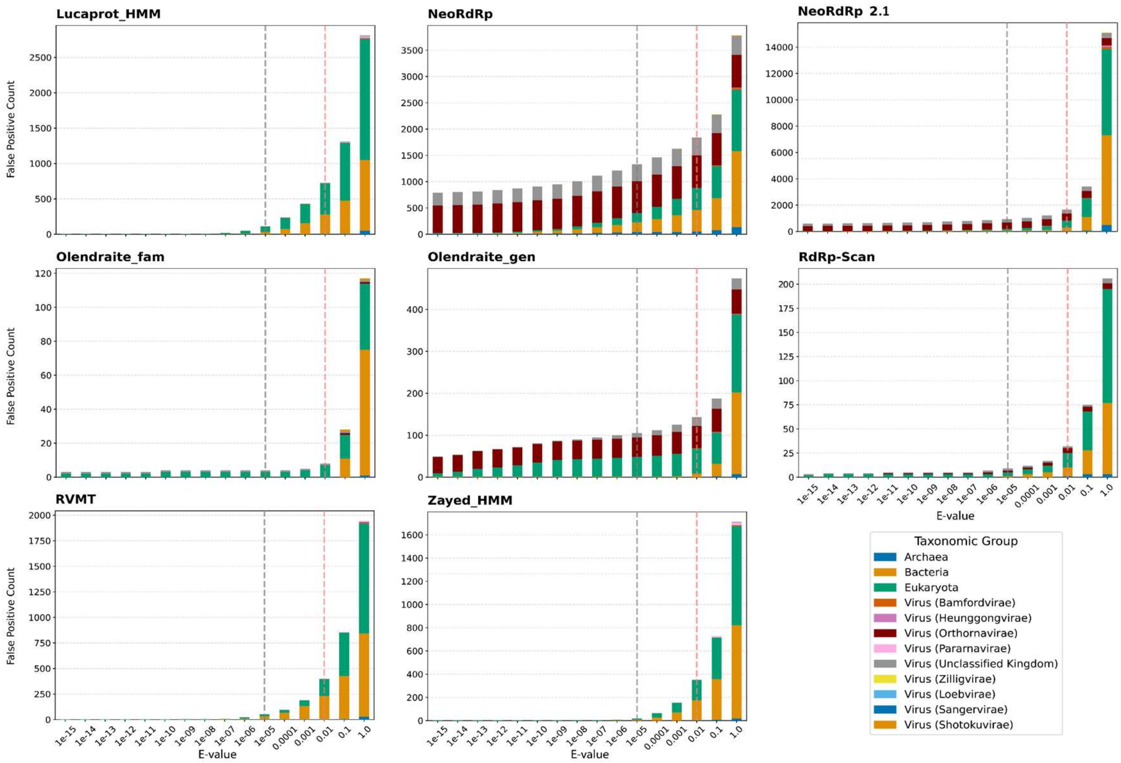

Figure S5. Taxonomic breakdown of false positive hits across E-value thresholds for comparative analysis scenario 2 (Total_TP/Total_TN dataset). Dashed lines indicate an E-value of $10^{-5}$ (grey) and $10^{-2}$(light red) to indicate suggested and default thresholds for hmmsearch.

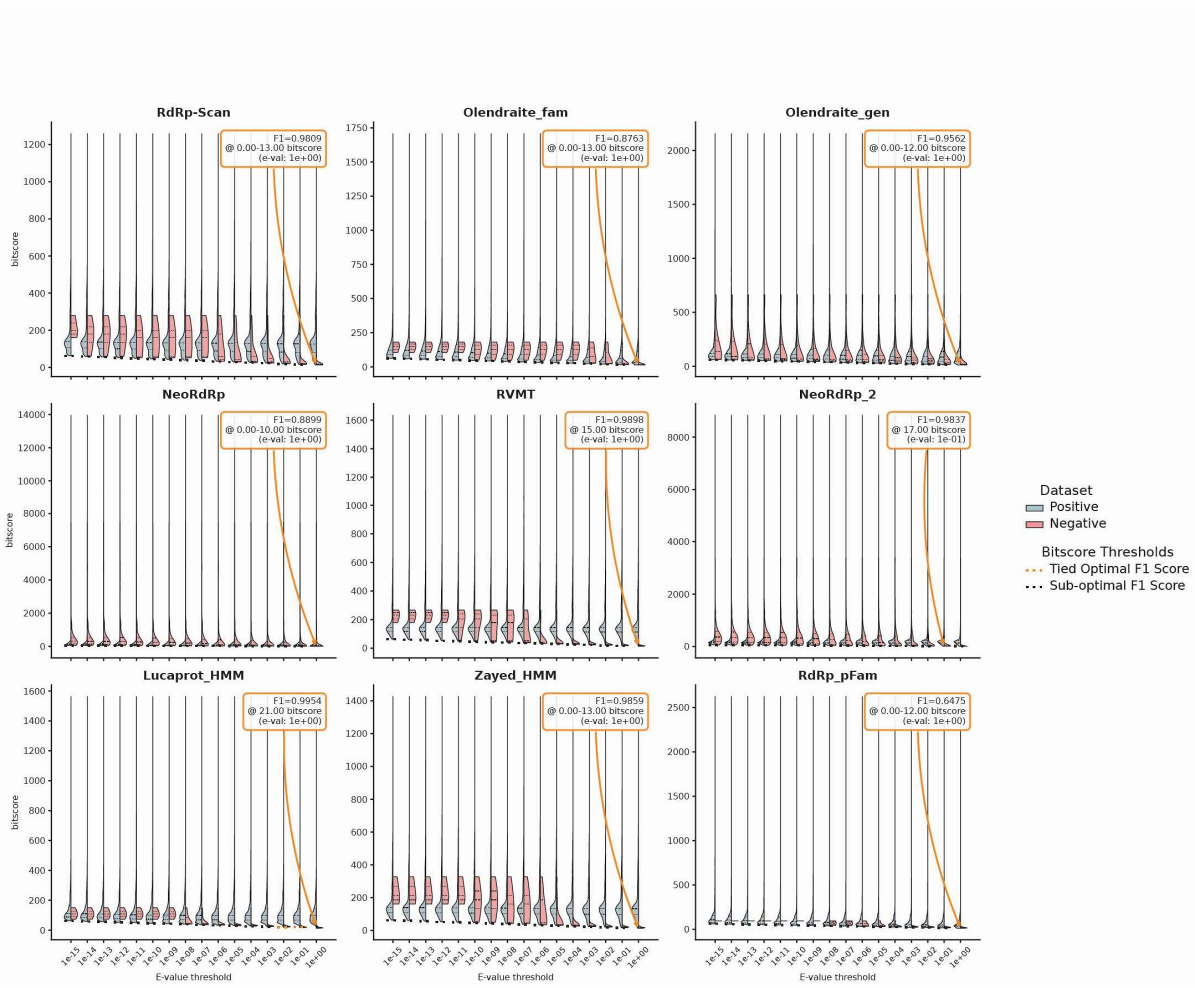

Figure S6. Bitscore distributions of true and false positive hits across different E-value thresholds and HMM databases for comparative analysis scenario 2 (Total_TP/Total_TN dataset). Each panel corresponds to a specific HMM database. For each E-value, the optimal bitscore threshold was determined by maximizing the F1-score. An annotation arrow highlights the overall best-performing E-value per database and its corresponding optimal bitscore, with the text box displaying the maximum F1-score achieved for that database. Dotted orange lines indicate bitscore thresholds for other E-values that also achieved this maximum F1-score (tie-break candidates), while dotted black lines show the optimal bitscores for the remaining, sub-optimal E-value thresholds.

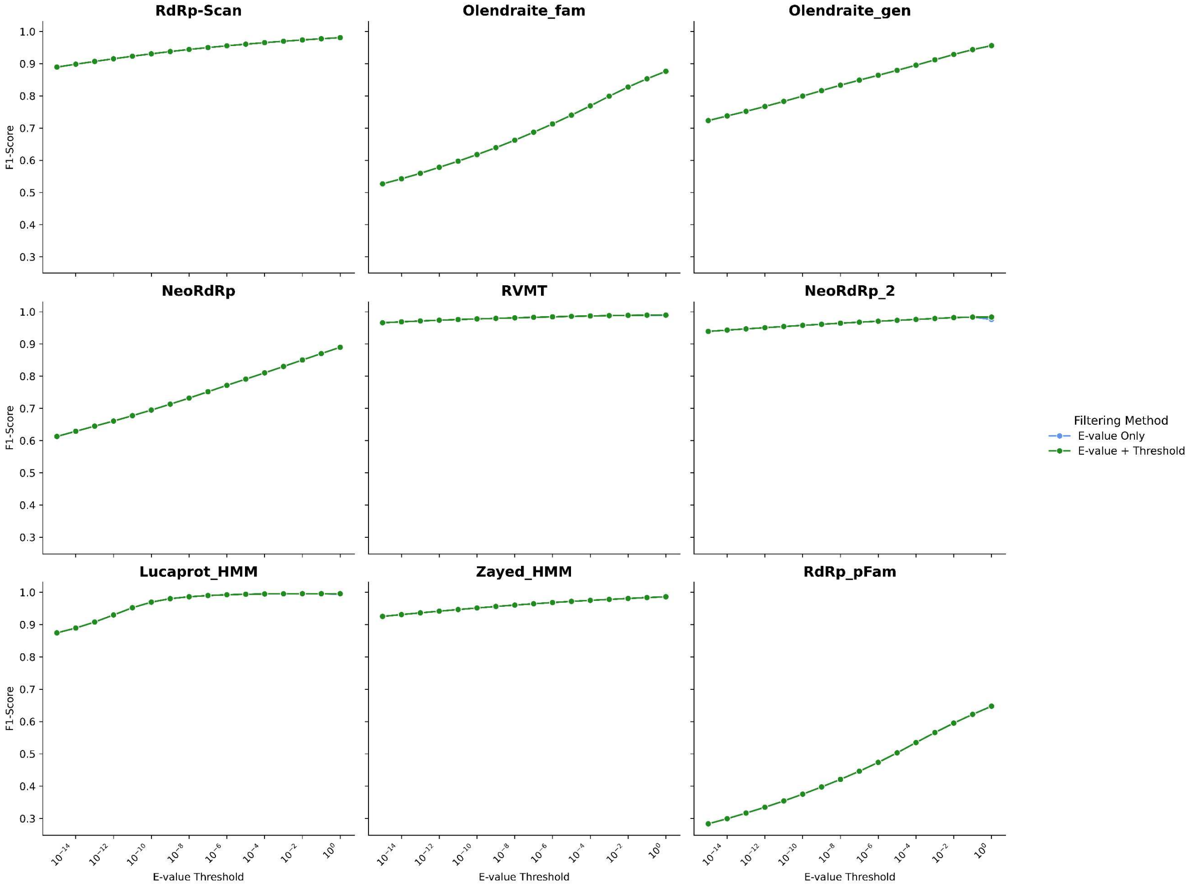

Figure S7.Improvement in F1-score after applying optimized bitscore filtering for comparative analysis scenario 2 (Total_TP/Total_TN dataset)

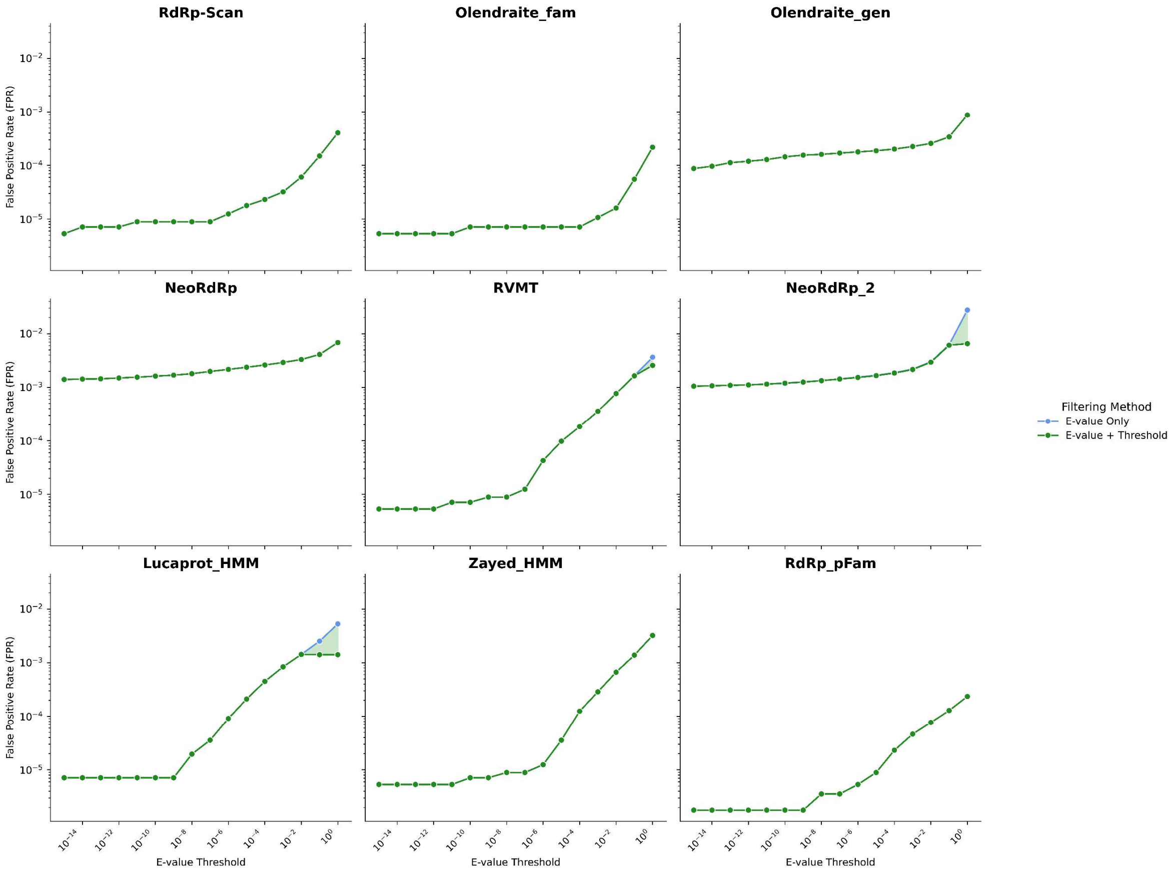

Figure S8. Reduction in False Positive Rate (FPR) after applying F1-optimized bitscore filtering for comparative analysis scenario 2 (Total_TP/Total_TN dataset).

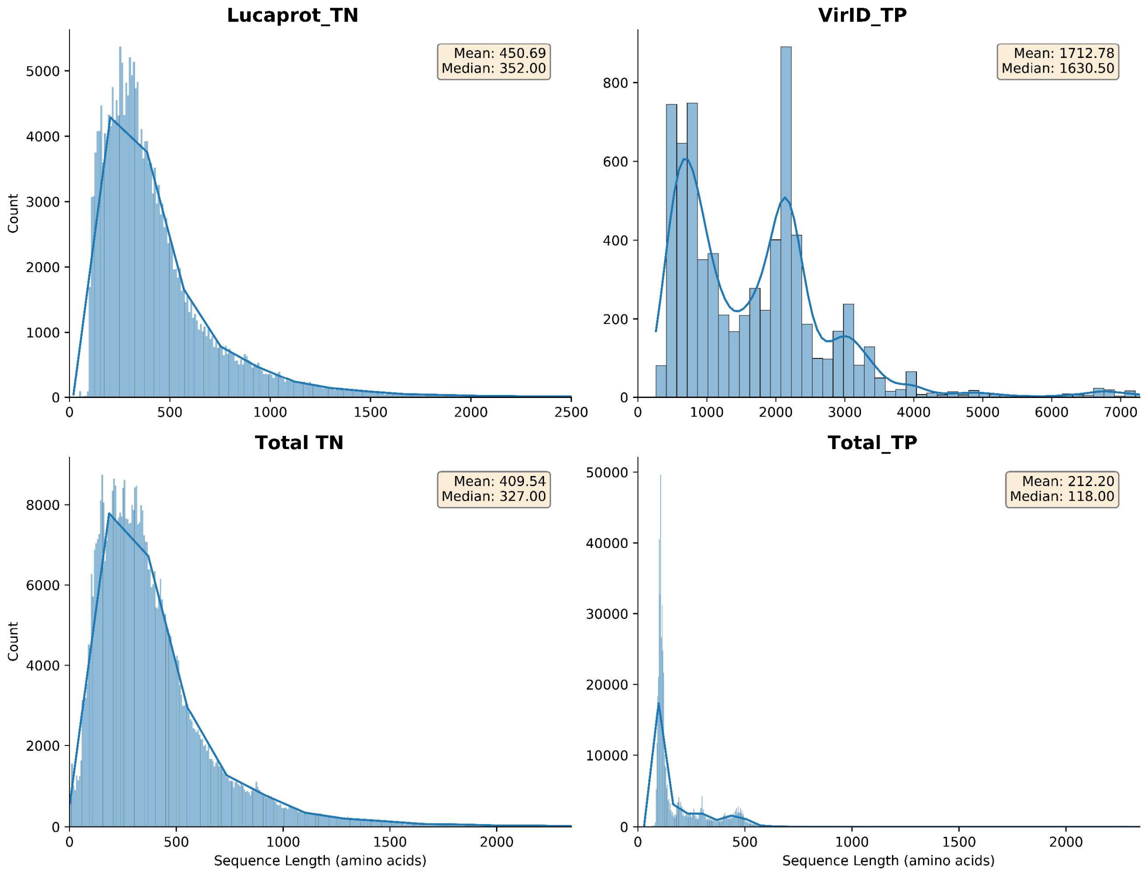

Figure S9. Protein sequence length distributions (in amino acids) for each of the four input FASTA files used in this study.

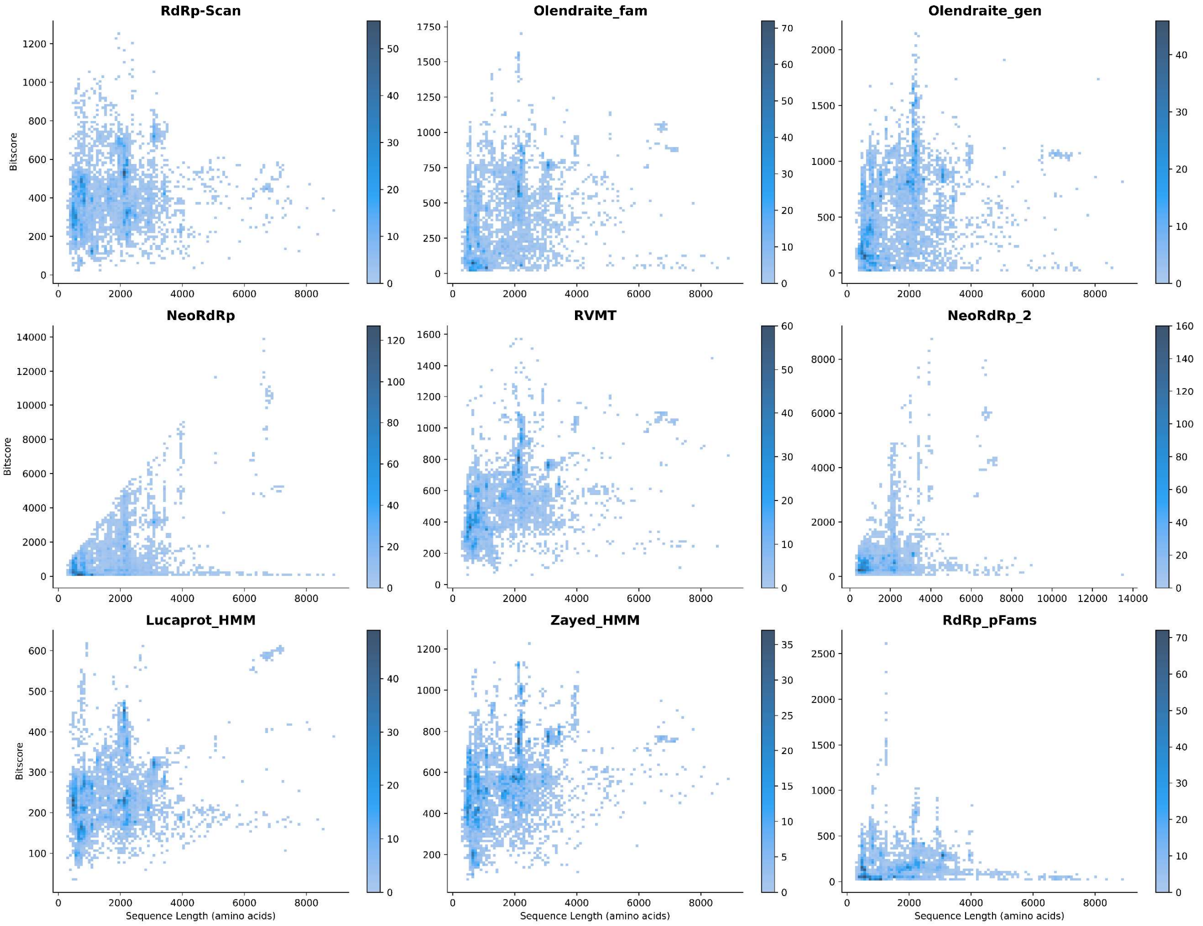

Figure S10. Density plot of HMM bitscore versus sequence length for true positive hits at a fixed E-value threshold of 1.0 for comparative analysis scenario 1 (VirID_TP/VirID_TN dataset).The general positive correlation observed across the databases indicates that longer sequences tend to produce higher bitscores, even at a permissive E-values.

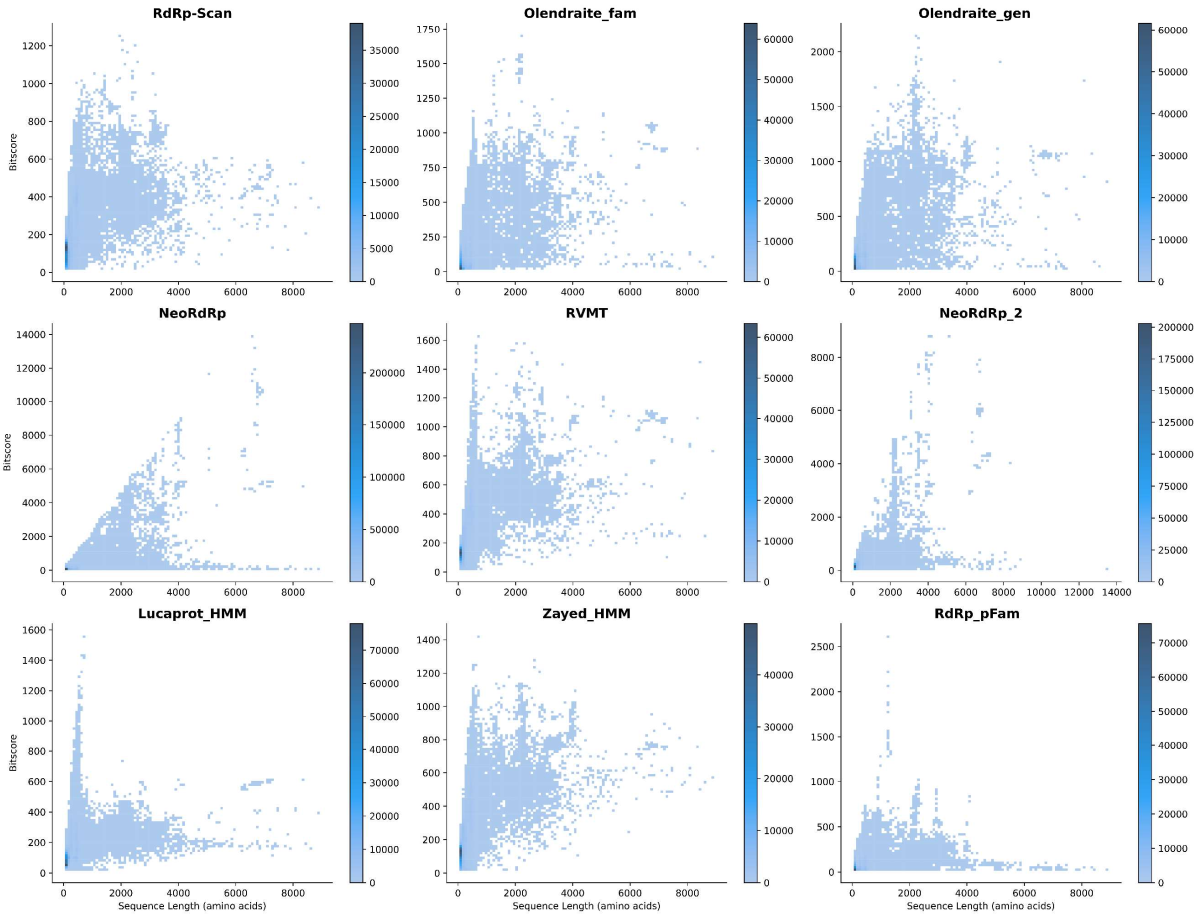

Figure S11. Density plot of HMM bitscore versus sequence length for true positive hits at a fixed E-value threshold of 1.0 for comparative analysis scenario 2 (Total_TP/Total_TN dataset). The general effect observed is that shorter sequences tend to have smaller bitscore values.

### References

1. Hou,X., He,Y., Fang,P., Mei,S.-Q., Xu,Z., Wu,W.-C., Tian,J.-H., Zhang,S., Zeng,Z.-Y., Gou,Q.-Y., *et al.* (2024) Using artificial intelligence to document the hidden RNA virosphere. *Cell*, **187**, 6929-6942.e16.

2. Sakaguchi,S., Urayama,S., Takaki,Y., Hirosuna,K., Wu,H., Suzuki,Y., Nunoura,T., Nakano,T. and Nakagawa,S. (2022) NeoRdRp: A Comprehensive Dataset for Identifying RNA-dependent RNA Polymerases of Various RNA Viruses from Metatranscriptomic Data. *Microbes Environ.*, **37**, ME22001.

3. Sakaguchi,S., Nakano,T. and Nakagawa,S. (2024) NeoRdRp2 with improved seed data, annotations, and scoring. *Front. Virol.*, **4**.

4. Olendraite,I., Brown,K. and Firth,A.E. (2023) Identification of RNA Virus–Derived RdRp Sequences in Publicly Available Transcriptomic Data Sets. *Mol. Biol. Evol.*, **40**, msad060.

5. Olendraite,I. (2021) Mining Diverse and Novel RNA Viruses in Transcriptomic Datasets . 10.17863/CAM.76470.

6. Charon,J., Buchmann,J.P., Sadiq,S. and Holmes,E.C. (2022) RdRp-scan: A bioinformatic resource to identify and annotate divergent RNA viruses in metagenomic sequence data. *Virus Evol.*, **8**, veac082.

7. Neri,U., Wolf,Y.I., Roux,S., Camargo,A.P., Lee,B., Kazlauskas,D., Chen,I.M., Ivanova,N., Allen,L.Z., Paez-Espino,D., *et al.* (2022) Expansion of the global RNA virome reveals diverse clades of bacteriophages. *Cell*, **185**, 4023-4037.e18.

8. Zayed,A.A., Wainaina,J.M., Dominguez-Huerta,G., Pelletier,E., Guo,J., Mohssen,M., Tian,F., Pratama,A.A., Bolduc,B., Zablocki,O., *et al.* (2022) Cryptic and abundant marine viruses at the evolutionary origins of Earth's RNA virome. *Science*, **376**, 156–162.

9. Babaian,A. and Edgar,R. (2022) Ribovirus classification by a polymerase barcode sequence. *PeerJ*, **10**, e14055.

10. Yang,Z., Shan,Y., Liu,X., Chen,G., Pan,Y., Gou,Q., Zou,J., Chang,Z., Zeng,Q., Yang,C., *et al.* (2024) VirID: Beyond Virus Discovery—An Integrated Platform for Comprehensive RNA Virus Characterization. *Mol. Biol. Evol.*, **41**, msae202.

11. Wolf,Y.I., Silas,S., Wang,Y., Wu,S., Bocek,M., Kazlauskas,D., Krupovic,M., Fire,A., Dolja,V.V. and Koonin,E.V. (2020) Doubling of the known set of RNA viruses by metagenomic analysis of an aquatic virome. *Nat. Microbiol.*, **5**, 1262–1270.

12. The UniProt Consortium (2025) UniProt: the Universal Protein Knowledgebase in 2025. *Nucleic Acids Res.*, **53**, D609–D617.
